# Transcription coordinates histone amounts and genome content

**DOI:** 10.1101/2020.08.28.272492

**Authors:** Kora-Lee Claude, Daniela Bureik, Petia Adarska, Abhyudai Singh, Kurt M. Schmoller

## Abstract

Biochemical reactions typically depend on the concentrations of the molecules involved, and cell survival therefore critically depends on the concentration of proteins. To maintain constant protein concentrations during cell growth, global mRNA and protein synthesis rates are tightly linked to cell volume. While such regulation is appropriate for most proteins, certain cellular structures do not scale with cell volume. The most striking example of this is the genomic DNA, which doubles during the cell cycle and increases with ploidy, but is independent of cell volume.

Here, we show that the amount of histone proteins is coupled to the DNA content, even though mRNA and protein synthesis globally increase with cell volume. As a consequence, and in contrast to the global trend, histone concentrations (i.e. amounts per volume) decrease with cell volume but increase with ploidy. We find that this distinct coordination of histone homeostasis and genome content is already achieved at the transcript level, and is an intrinsic property of histone promoters that does not require direct feedback mechanisms. Mathematical modelling and histone promoter truncations reveal a simple and generalizable mechanism to control the cell volume- and ploidy-dependence of a given gene through the balance of the initiation and elongation rates.

## Introduction

Maintaining accurate protein homeostasis despite cell growth and variability in cell volume is essential for cell function. Most proteins need to be kept at a constant, cell-volume-independent concentration. Since the amount of ribosomes and transcriptional machinery increases in proportion to cell volume, constant protein concentrations can be achieved through machinery-limited protein biogenesis, where protein synthesis depends on the availability of limiting machinery components and thus increases in direct proportion to cell volume^1,2^. While machinery-limited regulation can maintain constant concentrations of proteins, total mRNA, and individual transcripts^3–6^, it poses a conundrum for histones. As components of nucleosomes, histones are likely needed at a constant protein-to-DNA stoichiometry, implying that their amount should increase with ploidy but be independent of cell volume. In other words, histone concentration, *i.e*. amount per volume, should increase with ploidy but decrease with cell volume. Since accurate histone homeostasis is crucial for fundamental biological processes^7–10^ and to avoid toxic effects^11–13^, cells use several layers of regulation by translation, transcription and degradation to tightly coordinate histone production with genome replication^14–16^. However, how cells produce histones in proportion to genome content, even though protein biogenesis is generally linked to cell volume remains unclear.

Here, we use budding yeast as a model to show that histone protein amounts are coupled to genome content, resulting in a decrease of histone concentration in inverse proportion with cell volume, and an increase in direct proportion with ploidy. We find that this specific regulation of histones is achieved at the transcript level and does not require direct feedback mechanisms. While our data suggest that 3’-to-5’-degradation by the nuclear exosome is necessary for the correct decrease of concentration with cell volume, we show that histone promoters alone are sufficient to couple transcript amounts to gene copy number rather than cell volume. Our results suggest that this differential regulation of histones can be achieved through template-limited transcription, where mRNA synthesis is limited by the gene itself and does therefore not increase with cell volume. This provides a general mechanism by which cells can couple the amount of a subset of proteins to genome content while most protein concentrations are maintained constant.

## Results

### Histone protein concentrations decrease with cell volume and increase with ploidy

Typically, total protein amounts as well as the amounts of individual types of protein increase roughly in direct proportion to cell volume to maintain constant concentrations. However, such regulation is inappropriate for histones, whose amount we predicted should be coupled to the cellular genome content instead. To test if this is the case, we chose the budding yeast histone *HTB2*, one of two genes encoding for the core histone H2B, as an example, because it can be fluorescently tagged without pronounced effects on cell growth. We endogenously tagged *HTB2* with the fluorescent protein *mCitrine* in a haploid strain, and measured cell volume and the amount of Htb2-mCitrine as a function of time in cycling cells by microfluidics-based live-cell fluorescence microscopy^17,18^. To obtain a large range of cell volumes, we grew cells on synthetic complete media with 2% glycerol 1% ethanol as a carbon source (SCGE). As expected^14^, we find that Htb2 amounts are constant during early G1, rapidly double during S-phase and reach a plateau before cytokinesis (**Fig. 1a**). We then quantified the Htb2-mCitrine amounts in new-born cells directly after cytokinesis and find that the amount of Htb2-mCitrine is largely constant, independent of cell volume (**Fig. 1b**). To further test whether histone amounts are coupled to genomic DNA content rather than cell volume, we next analyzed a diploid strain in which both alleles of *HTB2* are tagged with *mCitrine*. Indeed, Htb2-mCitrine amounts in diploid cells are approximately a factor of two higher than in haploid cells (**Fig. 1b**). To more accurately compare Htb2 concentrations in haploids and diploids of similar volume, we sought to increase the overlapping range of observable volumes in both strains. For this purpose, we deleted the endogenous alleles of the G1/S inhibitor *WHI5* and integrated one copy of *WHI5* expressed from an artificial, β-estradiol-inducible promoter system^19^ (**Fig. 1c**). Using this system, we were able to increase the mean volume of steady-state exponentially growing populations by up to three-fold through overexpression of Whi5 (**Fig. 1d**) without drastically affecting doubling times, budding indices or cell cycle distributions (**Supplementary Fig. 1**). We repeated the microscopy experiments described above with the inducible-Whi5 haploid and diploid strains in the presence or absence of β-estradiol. Again, we find that Htb2-mCitrine amounts are only very weakly dependent on cell volume, but show a roughly two-fold increase in diploid compared to haploid cells (**Supplementary Fig. 2a**). Consistently, we find that the concentration of Htb2-mCitrine at birth in both haploid and diploid cells decreases strongly with cell volume (**Fig. 1e**). To quantify this decrease, we performed a linear fit to the double-logarithmic data, and defined the slope as the ‘volume-dependence-parameter’ (VDP). The observed VDPs of −0.87 ± 0.04 (haploids) and −0.97 ± 0.03 (diploids), respectively, are close to the value of −1 expected for proteins that are maintained at constant amount, resulting in a decrease of concentration with *c*~1/*V*. In contrast, proteins that are maintained at constant concentration would show a VDP of 0.

**Figure 1.**
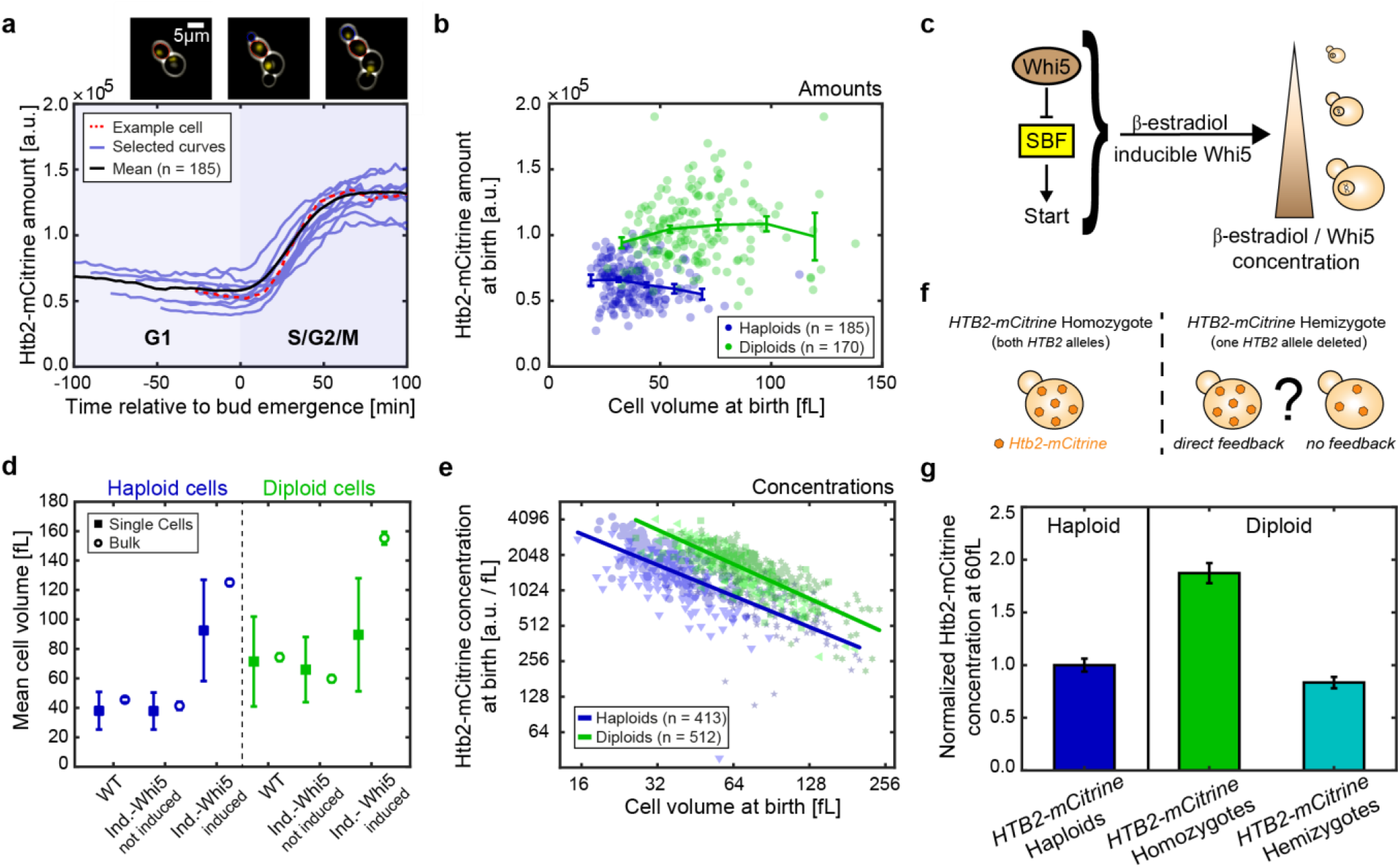
Htb2-mCitrine protein concentrations measured by live-cell fluorescence microscopy decrease with cell volume and increase with ploidy. (a) Htb2-mCitrine amounts during the first cell cycle of new-born cells. Red dashed trace highlights data corresponding to cells shown in the microscopy images (new-born cell: red outline, its bud: blue outline), blue traces show additional randomly selected example curves, black line the mean of n = 185 cells. All traces are aligned at the time of first bud emergence (t = 0). (b) Htb2-mCitrine amounts at birth for haploid (blue) and diploid (green) cells as a function of cell volume. Lines connect binned means, error bars indicate standard errors. (c) Whi5 controls cell volume in a dose-dependent manner. To manipulate cell volume, the endogenous allele is replaced by a copy of *WHI5* expressed from an artificial, β-estradiol-inducible promoter. Adding higher β-estradiol concentrations results in cells with bigger mean cell volumes. (d) Mean cell volumes for non-inducible (WT) and inducible haploids (blue) and diploids (green) measured in *HTB2-mCitrine* single cells with live-cell fluorescence microscopy (■), or in bulk populations of cells with untagged *HTB2* with a Coulter counter (○). Error bars indicate standard deviations of the mean between single cells for single cell measurements 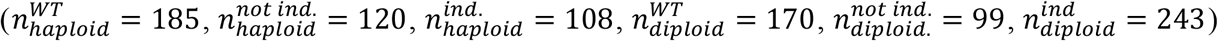 or the standard deviation of the population means across 5 biological replicates for bulk measurements. Haploid cells were induced with 30 nM β-estradiol, diploid cells with 50 nM. Note that no β-estradiol was used in the microfluidic device during the microscopy experiments, resulting in a gradual decrease of cell volume of induced cells after the start of the experiment. (e) Htb2-mCitrine concentrations of non-inducible and inducible haploids and diploids as a function of cell volume are shown in a double logarithmic plot. Individual data points for the different conditions (▼ 0 nM, ● WT, ★ 30 nM, for haploids and ◀ 0 nM, ■ WT, ✶ 50 nM, for diploids) are highlighted in blue (haploids) and green (diploids). Lines show linear fits to the double logarithmic data. (f) Illustration of the impact of potential feedback mechanisms on the concentration of Htb2-mCitrine concentration in a *HTB2-mCitrine/htb2Δ* hemizygous diploid compared to a *HTB2-mCitrine* homozygous diploid. (g) Htb2-mCitrine concentrations at 60 fL for haploids (blue), *HTB2-mCitrine* homozygous diploids (green) and *HTB2-mCitrine/htb2Δ* hemizygous diploids (teal) normalized on concentration at 60 fL in haploids. Error bars are derived by error propagation of the 95% confidence interval of the linear fit at 60 fL.

In budding yeast, histones are known to be tightly regulated at several layers. In particular, some histone genes – but not *HTB2* – exhibit dosage compensation at the transcript level^20–22^. In addition, excess histones are known to be degraded^16^. In principle, a coupling of histone amounts to genomic DNA content could be achieved through such feedback mechanisms: For example, larger cells may produce histones in excess, and then degrade the surplus. Alternatively, direct feedback of histone protein concentration on transcription could ensure that histones are expressed only until the protein amount matches the genome content. To test whether direct feedback of histone amounts on transcription, translation, or degradation is necessary to couple histone production to genome content, we again focused on Htb2, because it was already shown to not exhibit dosage compensation at the transcript level^21^. We constructed an inducible-Whi5 diploid strain in which we deleted one of the two *HTB2* alleles, while the other allele is tagged with *mCitrine* (**Fig. 1f**). If feedback were responsible for the coupling of Htb2 amount to genome content, the remaining *HTB2-mCitrine* allele should at least partially compensate for the deleted allele. However, consistent with the absence of any feedback, we find that Htb2-mCitrine concentrations are reduced by factor of two in the hemizygous compared to the homozygous diploid (**Fig. 1g, Supplementary Fig. 2b**). Moreover, at a characteristic volume of 60 fL, at which we find both haploid and diploid new-born cells, the concentration of Htb2-mCitrine in the hemizygous strain roughly equals the concentration in the haploid (**Fig. 1g**). While it is still possible that the reduced concentration of Htb2-mCitrine is compensated by an increased concentration of the other H2B copy Htb1, our results suggest that no direct feedback is required to couple Htb2 amounts to genome content. Instead, Htb2 amounts are intrinsically determined by the *HTB2* gene copy number, independent of ploidy and cell volume.

### Histone mRNA concentrations decrease with cell volume

The fact that the decrease of histone protein concentrations with cell volume is not simply a consequence of feedback, for example through excess protein degradation, suggests that it might already be established at the transcript level. To test if this is the case, we again employed the Whi5-overexpression system to measure the cell-volume-dependence of transcript concentrations (**Fig. 2a**). Specifically, we grew wild-type haploid cells, as well as the inducible-Whi5 haploid cells at three different β-estradiol concentrations (0, 10 and 30 nM), on SCGE media, which lead to a roughly four-fold range in mean cell volumes ranging from 39 ± 4 fL to 143 ±21 fL (**Supplementary Fig. 3a**). To ensure steady state conditions, we grew cells for at least 24 hours at the respective β-estradiol concentration, before then measuring cell volume distribution, extracting total RNA, and performing reverse-transcription-qPCR (RT-qPCR). First, we measured the concentration of the ribosomal RNA *RDN18* relative to total RNA and found it to be constant (**Supplementary Fig. 4a**). This is consistent with the fact that ribosomal RNA constitutes the large majority of total RNA^23^, which itself is expected to increase in direct proportion to cell volume^24^ and allows us to now normalize other RT-qPCR measurements on *RDN18*.

**Figure 2.**
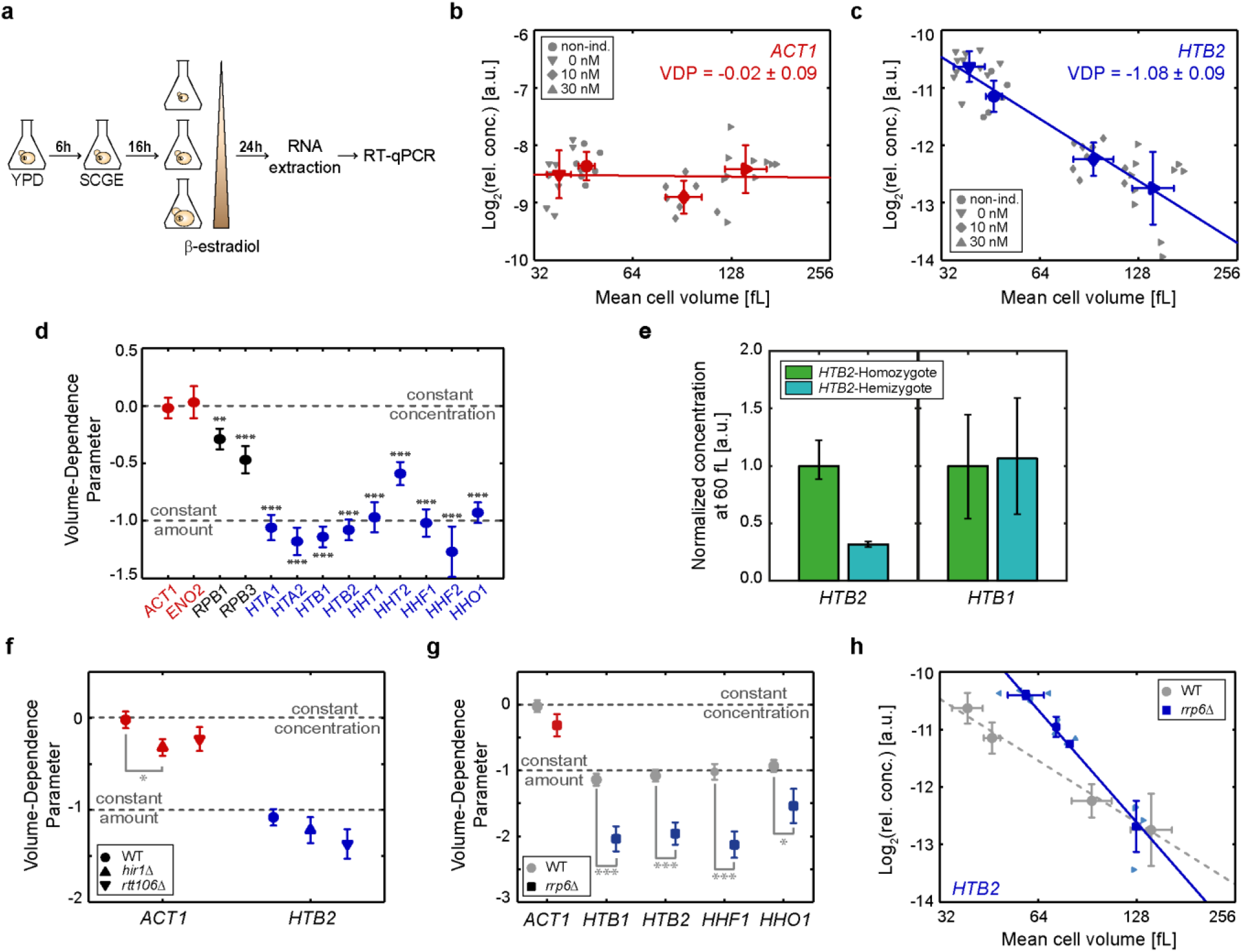
Histone mRNA concentrations decrease with cell volume and increase with ploidy. (a) Experimental procedure for RT-qPCR measurements. Cells were grown for at least 24 h at the respective β-estradiol concentration before extracting total RNA and performing RT-qPCR. (b & c) Relative *ACT1* (b) or *HTB2* (c) mRNA concentrations (normalized on *RDN18*) for non-inducible and inducible haploid cells over mean cell volume are shown in a double logarithmic plot. Individual data points for the different conditions (▼ 0 nM, ● non-inducible, ◆ 10 nM, ▲ 30 nM) are highlighted in grey. Red (b) or blue (c) symbols indicate the mean of the different conditions. Error bars indicate standard deviations for n ≥ 7 biological replicates. Lines show linear fits to the double logarithmic data, with volume-dependence parameters (VDPs) determined as the slope of the fit. (d) Summary of the VDPs for all measured genes. Error bars indicate the standard error of the slope; significances that the VDP is different from 0: **p<0.01, ***p<0.001. (e) Median mRNA concentrations at 60 fL of *HTB2* (left) and *HTB1* (right) in diploid *HTB2* homozygous (green) and *HTB2/htb2Δ* hemizygous (teal) strains, normalized on the respective median concentration of the *HTB2*-homozygote. Error bars indicate the 2.5- and 97.5-percentiles determined from 10000 bootstrap samples. (f & g) Summary of VDPs for *hir1Δ* and *rtt106Δ* (f) as well as *rrp6Δ* (g) deletion strains. Error bars indicate the standard error. Significant VDP deviation from the wild-type VDP (carrying no deletion) was tested using linear regressions; *p<0.05, ***p<0.001. (h) Relative *HTB2* mRNA concentrations (normalized on *RDN18*) for inducible and non-inducible haploid cells over mean cell volume, shown in a double logarithmic plot. Data corresponding to the *rrp6Δ* cells are highlighted in blue. Light blue symbols highlight the different conditions (◆ non-inducible, ◀ 0 nM, ▲ 10 nM, ▶ 30 nM). Dark blue symbols (■) indicate the mean for each condition. Grey symbols (●) indicate the mean for each condition of the wild-type (carrying no deletion). Lines show the linear fits to the double logarithmic data.

Next, we quantified the mRNA concentrations of *ACT1* and *ENO2*, two genes that we expect to be expressed in proportion to cell volume such that the mRNA concentration are maintained constant. Indeed, we find that the VDPs for both transcripts are not significantly different from 0 (**Fig. 2b & d, Supplementary Fig. 4b**). Interestingly, as previously suggested^25^ we observe a slight decrease in concentration for the transcripts of the RNA polymerase II subunits *RPB1* and *RPB3* with increasing cell volume (**Fig. 2d, Supplementary Fig. 4c**). We then quantified the concentrations of the transcripts of all core histone genes as well as the H1-like histone *HHO1*. In budding yeast, all core histone genes are present as two copies and expressed from bidirectional promoters controlling pairs of *H2A-H2B*^26^ or *H3-H4*^27^, respectively. Since the two copies of each core histone show high sequence similarity, we performed additional tests using deletion strains where possible to ensure qPCR primer specificity (**Supplementary Table 2**). We find that all histone transcripts show a significant decrease in concentration with cell volume, which is specific to the Whi5-dependent cell volume increase (**Supplementary Fig. 3b – d**). The histone mRNAs mostly exhibit VDPs close to −1 (**Fig. 2c & d, Supplementary Fig. 4d**). Thus, histone mRNA concentrations decrease with cell volume to ensure constant amounts – in contrast to global transcription, which increases with cell volume.

### Hir1-dependent feedback is not necessary for cell-volume-dependence of histone mRNA concentrations

The observation that histone transcript concentrations decrease with *c*~1/*V* suggests that, similar to histone protein amounts (**Fig. 1e**), also histone transcript amounts are determined by gene copy number. We therefore measured the concentrations of representative histone transcripts in inducible-Whi5 diploids homozygous or hemizygous for *HTB2*. Again, we find that all histones analyzed exhibit a VDP close to −1 (**Supplementary Fig. 5a**), and as observed for Htb2 protein concentrations (**Fig. 1f**), the concentration of *HTB2* transcripts at a characteristic volume of 60 fL is clearly reduced in hemizygous compared to homozygous diploids (**Fig. 2e**). Moreover, we do not observe a significant overexpression of *HTB1* to compensate for the reduced *HTB2* transcript concentration (**Fig. 2e**).

So far, we have shown that in diploid cells with only one *HTB2* allele, the concentrations of *HTB2* transcript and protein are reduced compared to wild-type diploid cells. This highlights the absence of direct feedback mechanisms sensing and controlling the concentration of Htb2 with cell volume. However, extensive previous studies have shown that the eight budding yeast core histone genes show remarkably different modes of regulation. Specifically, only the gene pair *HTA1-HTB1* is known to exhibit dosage compensation, which is absent for *HTA2-HTB2*^20–22^. Moreover, three out of four core histone gene pairs, not including *HTA2-HTB2*, show negative feedback regulation of transcript concentration upon replication stress^14,28^. This feedback regulation is thought to be mediated by the HIR complex and to be dependent on *HIR1* and *RTT106*^29–31^. Thus, to test if HIR-dependent sensing and feedback regulation of histone transcript concentration may also be responsible for the cell-volume-dependence of HIR-regulated histone genes, we measured the cell-volume-dependence of representative histone genes (*HTB1, HTB2, HHF1, HHO1*) in *hir1*Δ and *rtt106*Δ strains. Strikingly, we find that neither Hir1 nor Rtt106 are needed for the decrease of concentration with cell volume for any of the tested histone transcripts (**Fig. 2f, Supplementary Fig. 5b**).

### 3’-to-5’-degradation by the nuclear exosome is not necessary for cell-volume-dependence of histone mRNA concentrations

The fact that the correct dependence of histone transcript concentration on cell volume does not require direct feedback suggests that instead it is an intrinsic property of either transcription rate or mRNA degradation. To test if degradation from the 3’-end by the nuclear exosome is required, we analyzed the cell-volume-dependence of histone transcript concentrations in strains where we deleted *RRP6*, a component of the nuclear exosome exonuclease^32,33^. As shown in **Fig. 2g**, we find that also in *rrp6*Δ cells, histone transcript concentrations decrease with cell volume. Interestingly, due to increased transcript concentrations in small cells (**Fig. 2h, Supplementary Fig. 5c**), this decrease with a VDP close to −2 is significantly stronger than in wild-type cells, suggesting that the volume-dependence of histone transcripts is modulated by Rrp6-dependent degradation. Thus, while degradation by the nuclear exosome is not needed for the volume-dependent decrease of histone transcript concentrations, it may contribute to achieve the correct VDP of −1.

### Histone promoters are sufficient for cell-volume-dependence of transcript concentrations

Given that degradation from the 3’-end does not seem to be crucial for the cell-volume-dependent decrease of histone transcript concentration, we next asked whether the promoter alone is sufficient to establish this cell-volume-dependence. To address this, we created strains that carry additional copies of either the *ACT1* or the histone *HHF1* promoter driving expression of the fluorescent protein mCitrine, regulated by the identical *ADH1* terminator (**Fig. 3a**). Strikingly, we find that the dependence of *mCitrine* transcript concentration on cell volume is determined by the promoter: If driven by the *ACT1* promoter, the VDP of *mCitrine* resembles that of endogenous *ACT1*; if driven by the *HHF1* promoter, it resembles that of endogenous *HHF1* (**Fig. 3b**).

**Figure 3.**
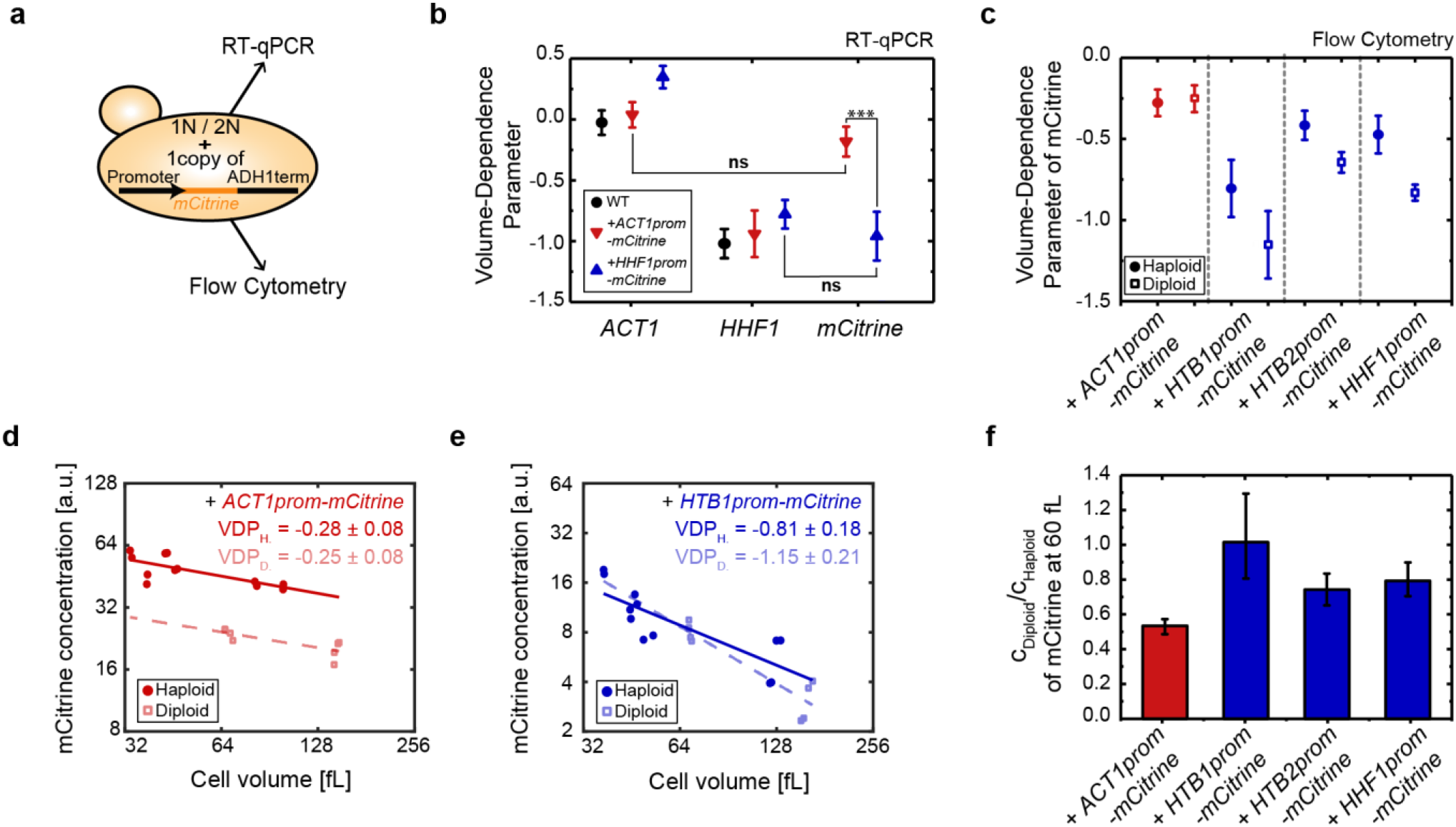
Histone promoters are sufficient for cell-volume- and ploidy-dependence of transcript concentrations. (a) Illustration of haploid (1N) or diploid (2N) strains carrying a single additional copy of a promoter of interest, driving the expression of the fluorescent reporter *mCitrine* regulated by the *ADH1* terminator. RT-qPCR or flow cytometry were used to analyze expression of the fluorescent reporter. (b) Summary of VDPs determined with RT-qPCR for the genes *ACT1, HHF1* and *mCitrine* in a wild-type strain (black •), a strain carrying an additional *ACT1* promoter (red ▼), or a strain carrying an additional *HHF1* promoter (blue ▲). Error bars indicate the standard error. Significant VDP deviation between two genes was tested using linear regressions; ***p<0.001. (c) Summary of VDPs determined with flow cytometry for different strains in haploid (●) and diploid (□) cells. Error bars indicate the standard error. (d – e) *mCitrine* concentration, driven by an additional copy of the *ACT1* (d) or *HTB1* (e) promoter in haploid (●) and diploid (□) cells, shown as a function of cell volume in a double logarithmic plot. Lines show linear fits to the double logarithmic data with volume-dependence parameters (VDPs) determined as the slope of the fit, with respective standard error. (f) Median concentration of mCitrine in diploid cells compared to the median concentration in haploid cells at 60 fL. Error bars indicate the 2.5- and 97.5-percentiles determined from 10000 bootstrap samples.

To test if this also holds true for other histone promoters, we made use of the fact that the fluorescent reporter mCitrine enables a faster experimental readout using flow cytometry (**Fig. 3a**). First, we analyzed the cell-volume-dependent fluorescence of mCitrine expressed from the *ACT1* or *HHF1* promoters, which revealed that flow cytometry can be used to qualitatively distinguish the distinct volume-dependences. Similarly, we find that also all other histone promoters tested show significantly negative VDPs in haploid and diploid cells (**Fig. 3c – e, Supplementary Fig. 6**).

Histones not only need to be maintained at cell-volume-independent amounts, leading to a decrease of concentration with 1/*V*, but also need to increase in proportion to cell ploidy (**Fig. 1**). This is in contrast to most other genes, which are maintained at a ploidy-independent concentration^34^. To test if the histone promoters are also sufficient to establish this distinct ploidy-dependence, we compared the expression level of the single *mCitrine* copy in diploid versus haploid cells. For *ACT1*, which needs to be maintained at a ploidy-independent concentration, we expect that a single gene allele in a diploid should produce half of the protein compared to a homozygous diploid or haploid of similar volume^2^. Indeed, for the *ACT1* promoter we find that at a given cell volume, the concentration of mCitrine expressed from a single additional promoter is 50% lower in diploids compared to haploids (**Fig. 3d & f**). In contrast, for each of the three histone promoters tested, we observe that the concentration in diploids is considerably higher than 50% of that in haploids of comparable volume, with a ratio close to 1 for the *HTB1* promoter (**Fig. 3e & f, Supplementary Fig. 6**). This demonstrates that in addition to setting the cell-volume-dependent decrease in concentration, regulation by the histone promoters also largely accounts for the fact that histones are needed in proportion to ploidy.

### Different cell-volume and ploidy dependences can be explained by competition of promoters for limiting transcriptional machinery

To better understand how the transcription rate of one specific promoter depends on cell volume and ploidy context, we sought to build a minimal model (**Fig. 4a**). Briefly, we considered two classes of promoters, a specific promoter of interest, *p*, present as a single copy, and a general pool of promoters, *g*, which are present as *n_h_* = 6000 in haploids or *n_d_* = 12000 copies in diploids. We then assume that transcription can be described by a single component of the transcriptional machinery, whose concentration *c_TM_* stays constant with cell volume. Each promoter is competing for the transcriptional machinery, and is modelled as a single binding site for the limiting machinery component. Initiation, *i.e*. binding of the limiting machinery, occurs at a rate 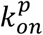 or 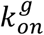, respectively. Furthermore, we assume that all other steps of transcription can be summarized in a single rate-limiting step, occurring at a rate 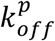 or 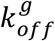, respectively. Each transcript is then degraded with the same rate *k_deg_* = 1. Depending on the parameters chosen for the specific promoter, the model predicts qualitatively different dependences of transcript concentration on cell-volume and ploidy (**Fig. 4b & c**). For example, at a given 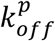, a high on-rate 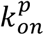 can result in histone-promoter-like behavior, *i.e*. cell volume-dependent but ploidy-independent transcript concentration. In contrast, at lower 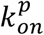 we observe actin-promoter-like behavior, *i.e*. cell volume-independent but ploidy-dependent transcript concentration. Interestingly, due to the competition with general promoters, the transcript concentration can even increase with cell volume if 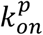 is much smaller than 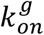.

**Figure 4.**
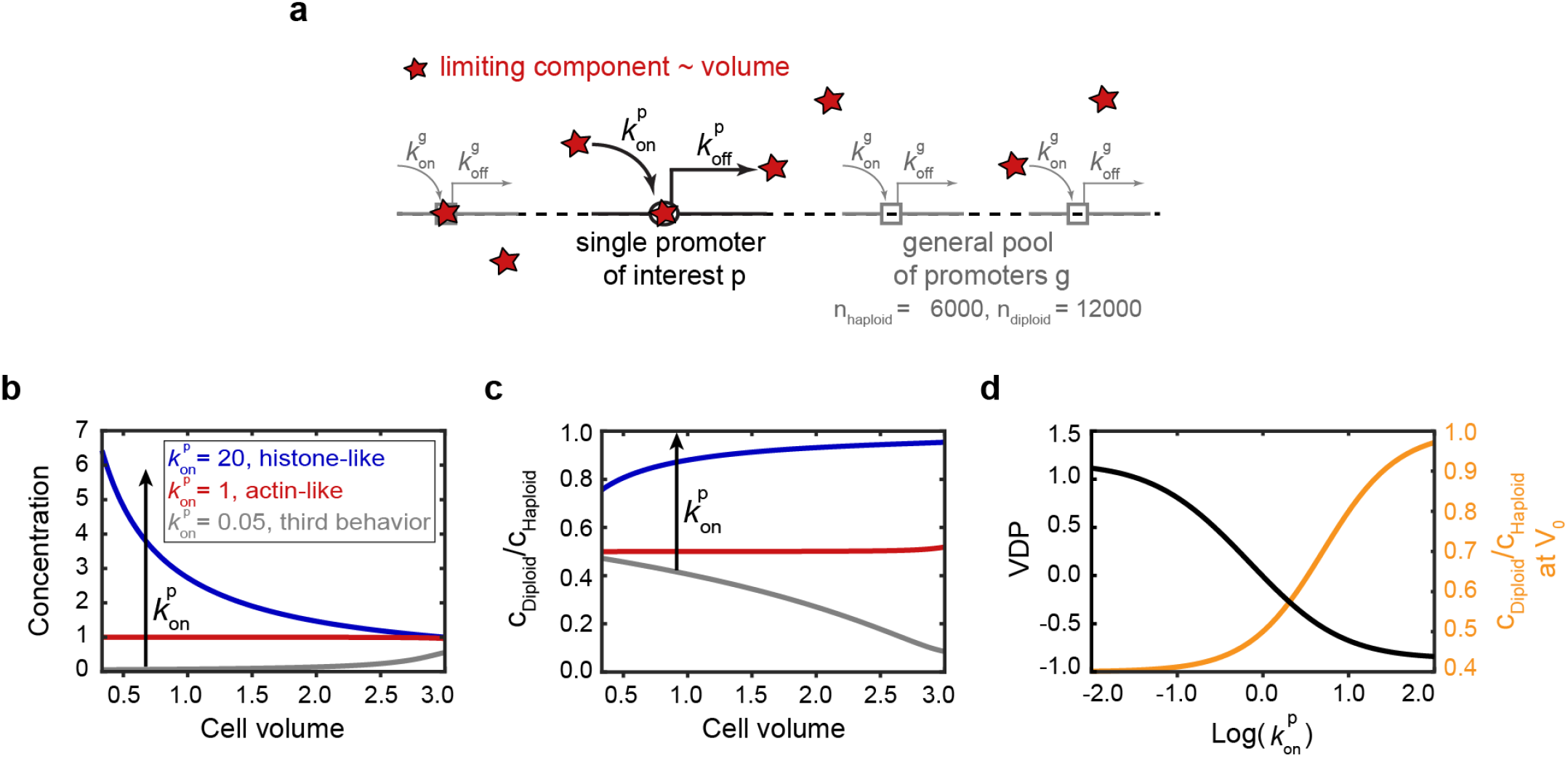
Minimal model for the dependence of transcription rate of one specific promoter of interest on cell volume and ploidy. (a) The model includes two classes of promoters: the general pool of promoters *g* and the specific promoter of interest p with their respective initiation rates 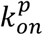 or 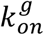, describing the binding of the limiting machinery and off-rates 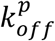 or 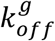, summarizing all other steps of transcription. (b - d) The model predicts that tuning 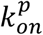 while keeping all other parameters fixed 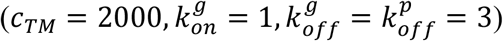 results in a qualitative change of the cell volume-dependence of transcript concentration obtained from the specific promoter (b), as well as a change in the ratio between the concentration in diploid cells and the concentration in haploid cells (c). (d) Model prediction for the VDP (right, black) and the ratio between the concentration in diploid cells and the concentration in haploid cells at a characteristic volume *V*_0_ = 1 (left, orange) as a function of 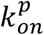.

One key prediction of this model is that if all other parameters are fixed, reducing 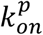 for a histone-like promoter should eventually shift its behavior to that of an actin-like promoter (**Fig. 4d**). To experimentally test this prediction, we aimed to decrease the initiation rate 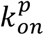 of the *HHF1* and *HTB1* promoters by creating series of haploid and diploid strains with increasingly shorter fragments of the promoters, each truncated from the 5’-end (**Fig 5a**). Again, we used flow cytometry to analyze mCitrine expression driven by these additional, endogenously integrated promoter fragments. For both promoters we observe a decrease of mCitrine expression once part of the known upstream activating sequences (UASs) are truncated (**Fig 5b, Supplementary Fig. 7a**). Fully consistent with the model, for both promoters, and for haploids and diploids, this drop in expression coincides with a change of the VDP towards 0 (**Fig 5b & c, Supplementary Fig. 7b & c**). At the same time and also consistent with the model, the ratio of the mCitrine concentration at a given volume in diploid compared to haploid cells decreases from close to 1 towards 0.5 (**Fig. 5c**). Thus, our analysis shows that for both the *HHF1* and *HTB1* promoter truncation series, a transition from histone-like to actin-like behavior occurs between the 450 bp to 300 bp truncations.

**Figure 5.**
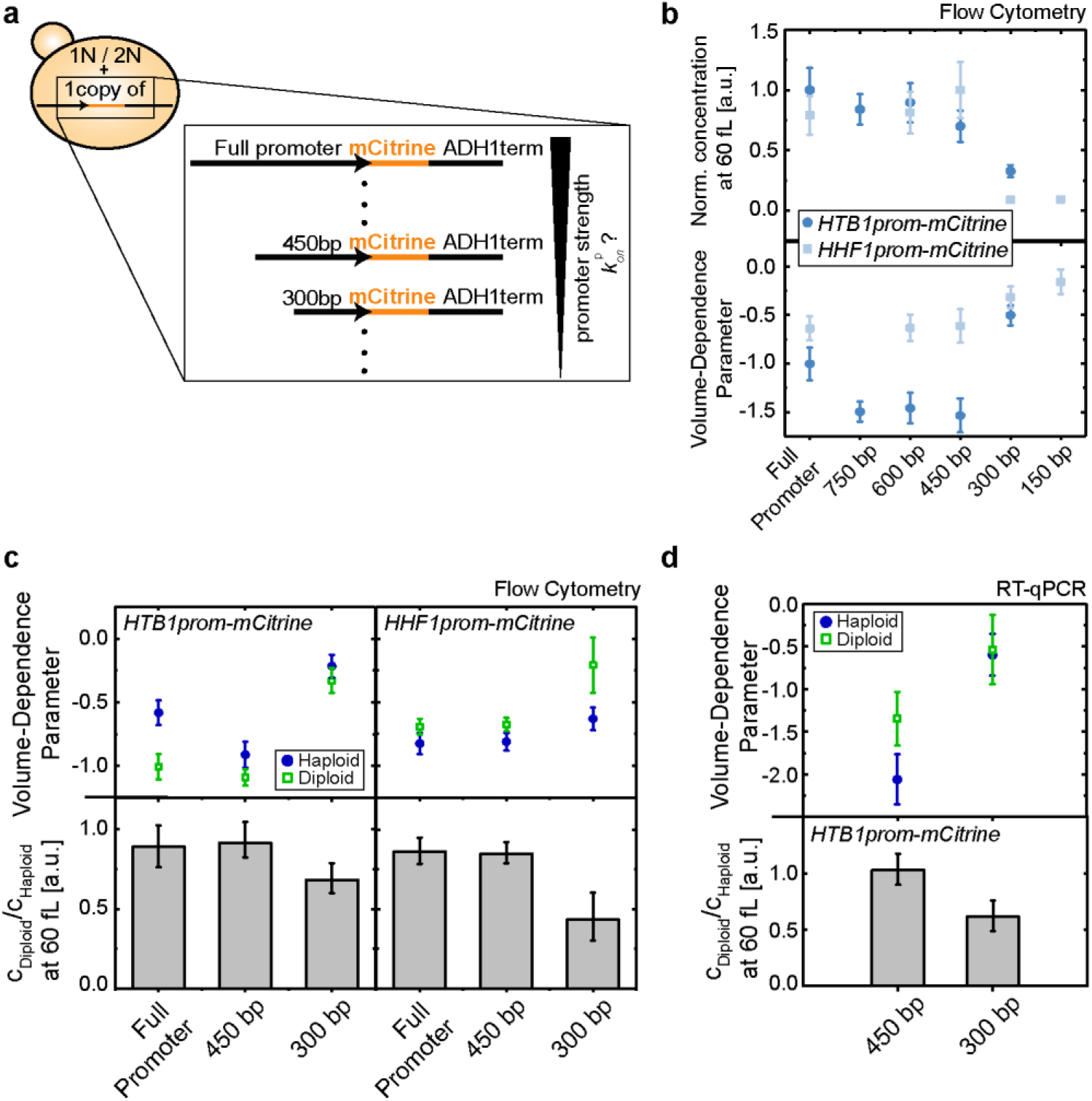
Reducing the strength of a histone promoter shifts its behavior from histone-like to actin-like. (a) Illustration of a series of haploid and diploid strains carrying a single additional copy of increasingly shorter fragments of promoters driving *mCitrine* expression, each truncated from the 5’-end. (b) mCitrine concentration at 60 fL normalized on maximum concentration of the respective promoter (upper panel) and VDP of mCitrine (bottom panel) determined by flow cytometry for the respective promoter truncations of the *HTB1* promoter (dark blue ●) and the *HHF1* promoter (light blue ■) driving *mCitrine* expression, integrated in haploid cells. Error bars in the upper panel are derived by error propagation of the 95% confidence interval of the linear fit at 60 fL. In the bottom panel, error bars show the standard error. (c) VDP of mCitrine in haploid (blue ●) and diploid (green □) cells (upper panel) and mCitrine concentration at 60 fL in diploids compared to the concentration in haploids (bottom panel) determined by flow cytometry. Left shows results for the *HTB1* promoter truncations, right shows results for the *HHF1* promoter truncations. Error bars in the upper panels show the standard error. In the bottom panel, error bars indicate the 2.5- and 97.5-percentiles determined from 10000 bootstrap samples. (d) VDP of *mCitrine* in haploid (blue ●) and diploid (green □) cells (upper panel) and *mCitrine* mRNA concentration at 60 fL in diploids compared to the concentration in haploids (bottom panel) determined by RT-qPCR for *HTB1* promoter truncations driving *mCitrine* expression. Error bars in the upper panel show the standard error. Error bars in the bottom panel indicate the 2.5- and 97.5-percentiles determined from 10000 bootstrap samples.

While we consistently observe the same qualitative trend in flow cytometry measurements, we note that the exact VDP depends on the forward scatter settings, which determine the observed cell-volume range. Thus, to quantitatively confirm our results, we repeated the experiment for the 450 bp and 300 bp truncations of the *HTB1* promoter using RT-qPCR. Again, we observe a change in the VDP towards 0, and a decrease of the ratio of the mCitrine concentration between diploid and haploid cells from close to 1 to close to 0.5 (**Fig. 5d**). In summary, our analysis of the histone promoter truncations demonstrates that decreasing promoter strength can shift the volume- and ploidy-dependence of the histone promoters to an actin-like behavior, as predicted by our minimal model.

## Discussion

Taken together, we identified a mechanism that allows cells to deal with a fundamental challenge – how to quantitatively couple histone production to DNA content even though total biosynthetic capacity is linked to cell volume instead. We found that this coordination is already achieved at the transcript level. While mRNA degradation and feedback mechanisms contribute to histone homeostasis, we find that competition for potentially limiting transcriptional machinery is sufficient to achieve differential regulation of histone and other transcript concentrations with cell volume and ploidy. Specifically, if transcription is limited by the availability of limiting machinery, larger cells with more machinery will produce proportionally more mRNA, maintaining constant transcript concentrations, which do not depend on ploidy. If transcription is instead limited by the gene itself, transcript concentrations will decrease with cell volume but will be proportional to ploidy. In addition to histones, other proteins will require differential regulation. For example, the G1/S inhibitors Whi5 in yeast^18^ and Rb in mammalian cells^36^ have recently been shown to decrease in concentration with cell volume, enabling cells to sense and control their size. Along those lines, a recent study suggested that many cell cycle regulators show differential transcriptional regulation with cell volume^37^. The simplicity of template-limited transcription therefore suggests that it may be broadly employed across species to differentially regulate the concentrations of larger subsets of proteins, in particular to couple the amount of DNA binding proteins to DNA content. Moreover, in addition to the ideal template- or machinery-limited regimes, cells can achieve a large variety of cell volume- and ploidy-dependences, which importantly can be decoupled from the expression level of a given gene by independently tuning its initiation and elongation rates. Specific regulation of mRNA and protein degradation provides yet another level of control that cells can employ to tune the dependence of protein concentrations on cell volume and ploidy. In fact, our observation that the cell-volume-dependence of histone transcripts is even stronger in *rrp6* deletion cells, suggests that such additional regulation contributes to cell-volume-dependent histone homeostasis in budding yeast. To quantitatively understand the cell volume- and ploidy-dependence of protein homeostasis on a genome wide level, it will therefore be crucial to identify the rate-limiting steps of transcription and mRNA degradation as well as the corresponding rate-limiting molecules.

## Materials and methods

### Yeast strains

All yeast strains used in this work are based on W303 and were constructed using standard methods. Full genotypes of all strains are listed in **Supplementary Table 1**.

### Inducible-Whi5 strain

In order to increase the range of observable cell volumes, we used strains with β-estradiol inducible *WHI5*, similarly described in previous works^18,38^. For this purpose, we deleted the endogenous alleles of the G1/S inhibitor *WHI5* and integrated one copy of *WHI5* expressed from an artificial, β-estradiol-inducible promoter system^19^. Specifically, this inducible promoter system consists of a β-estradiol-dependent, artificial transcription factor, which can bind an artificial promoter. This promoter is then used to induce *WHI5* expression.

To ensure that β-estradiol addition itself has no effect on cell growth, we grew cell cultures of a non-inducible *WHI5* haploid strain and cell cultures of a *whi5Δ* haploid strain, containing the β-estradiol-dependent, artificial transcription factor, but no copy of *WHI5*. We then added β-estradiol to those cultures and quantified the mean cell volumes after 24 h of growth in the presence of β-estradiol, by measuring the cell volume distributions using a Coulter Counter (Beckman Coulter, Z2 Particle Counter). Finally, we compared the mean cell volumes to the mean cell volumes obtained from cell populations without β-estradiol addition (**Supplementary Fig. 3a**). In addition, we performed reverse-transcription-qPCR (RT-qPCR) on cell populations with and without β-estradiol addition and compared the obtained mean values for several genes (**Supplementary Fig. 3b & c**). For the non-inducible *WHI5* haploid strain, we could not identify a significant deviation of the population means between the cell populations with and without β-estradiol addition. For the *whi5Δ* haploid strain, containing only the β-estradiol-dependent, artificial transcription factor, we observed a slight but significant reduction of the relative mean mRNA concentrations of *HTA2, HHF2* and *HHO1* at 30 nM compared to 0 nM β-estradiol, which was consistent with a slightly increased mean cell volumes at 30 nM β-estradiol. In contrast, performing the same experimental procedure on cell cultures of an inducible *WHI5* haploid strain, leads to much stronger changes of mean cell volumes and relative mean mRNA concentrations for all histone genes, demonstrating that the observed decrease of histone mRNA concentrations is specific to the Whi5-dependent cell volume increase (**Supplementary Fig. 3a & d**). Significances were tested using two-tailed two-sample t-tests, after checking for normal distribution and equal variance distributions using a Shapiro-Wilk test and a Bartlett test, respectively.

### Live-cell fluorescence microscopy

Cultures (3 mL) were grown at 30°C in synthetic complete media containing 2% glycerol and 1% ethanol (SCGE) for at least 6 h in a shaking incubator at 250 rpm (Infors, Ecotron). Appropriate β-estradiol concentrations were then added to inducible cells (0 nM and 30 nM for haploids or 50 nM for diploids) and the cultures grown for at least 24 h to ensure steady-state conditions. Optical densities were measured using a spectrophotometer (Perkin Elmer, Lambda Bio+) and *OD*_600_ < 1.0 was maintained through appropriate dilutions during culture growth. For imaging, 1 mL of cells (*OD*_600_ < 1.0) was spun down at 10k g-force for 1 min (Thermo Fisher Scientific, Pico 17), resuspended in 200 μL SCGE and sonicated for 5 s (Bandelin electronics, HD2070 & UW2070). 100 μL of this cell suspension was then introduced in a Cellasic microfluidics Y04C (haploids and non-induced diploids) or Y04D (induced diploids) plate.

Live-cell fluorescence microscopy experiments were performed on a Zeiss LSM 800 microscope with additional epifluorescence setup using a Cellasic microfluidics device to ensure constant media (SCGE) flow in the microfluidics plate throughout the experiment. Experiments ran for 12 h with images being taken every 3 min using an automated stage (WSB Piezo Drive Can), a plan-apochromat 40x/1.3 oil immersion objective and an axiocam 506 camera. Phase-contrast images were taken at an illumination voltage of 4.5 V and an exposure time of 30 ms. *mCitrine* images were taken using the Colibri 511 LED module at 25% power and an exposure time of 10 ms. For each condition, at least two independent biological replicates were measured on different days.

To correct for inaccuracies of the x-y-stage between time points, movies were first aligned using a custom Fiji script. Then, cell segmentation and quantification of the fluorescent signal as well as subtraction of background fluorescence and cell-volume-dependent autofluorescence (determined from control strains not expressing a fluorescent protein), and determination of time points of cell birth, bud emergence, and cytokinesis were performed with MATLAB 2017b using previously described methods^17,18,39^. For our analyses, we only included cells born during the experiment. Total fluorescence intensity after background- and autofluorescence correction was used as a proxy for total protein amount.

In order to determine total protein concentrations as total protein amounts divided by cell volume, we calculated cell volumes based on phase-contrast images. Briefly, after segmentation, cell areas where aligned along their major axis. We then divided the cells into slices perpendicular to their major axis, each 1 pixel in width. To estimate cell volume, we then assumed rotational symmetry of each slice around its middle axis parallel to the cell’s major axis, and summed the volumes of each slice to obtain total cell volume. This allowed us to analyze protein amounts and protein concentrations as a function of cell volume.

### Estimation of cell cycle phases and histone production period using live-cell microscopy

To test whether the decrease of histone concentrations with cell volume could be explained by a decrease in the S-phase duration, and thus a shorter time period during which histones are produced, we aimed to estimate the duration of the *histone production period* (H-period) from the Htb2-mCitrine fluorescent intensity traces. For each single cell, we first performed a constant linear fit in each of the two plateaus of the fluorescence intensity, linked to G1- or G2/M-phase, respectively, and denoted them as *P*_1_ and *P*_2_. *P*_1_ was obtained by performing the linear fit through the data points of the fluorescent intensity trace from cell birth to first bud emergence, *P*_2_ was obtained by performing the linear fit through the last 30 minutes of the fluorescent intensity trace. We then set a threshold of 5%, determined the last time point for which *I_Htb2–mCitrine_* < *P*_1_ + 0.05 · *P*_1_, and defined this time point as the beginning of the H-period. Similarly, we defined the first time point for which *I_Htb2-mCitrine_* > *P*_2_ – 0.05 · *P*_2_ as the end of the H-period. Finally, the duration of the H-period was calculated as the difference between those two time points. We defined G1-phase duration as the time from cell birth to first bud emergence, and G2/M duration as the time between the end of the H-period and cytokinesis.

### RNA extraction and RT-qPCR

Cultures (25 mL) were grown at 30°C in yeast peptone media containing 2% glucose (YPD) for at least 6 h in a shaking incubator at 250 rpm, before being washed and transferred to SCGE. The cultures were grown for at least 16 h before appropriate β-estradiol concentrations were added to inducible cells (0 nM, 10 nM and 30 nM). The cultures (final volume of 50 mL) were then grown for at least 24 h in order to ensure steady-state conditions. During culture growth, *OD*_600_ < 1.0 was maintained through appropriate dilutions. Cell volume distributions of the cultures were measured with a Coulter counter after sonication for 5 s.

Remaining cell cultures were spun down at 4000 rpm for 5 min and the cell pellet resuspended in 50 μL nuclease-free water (Qiagen). Total RNA was extracted using a hot acidic phenol (Sigma-Aldrich) and chloroform (Thermo Fisher Scientific) extraction method adapted from an established protocol^40^. Yield of RNA was increased by precipitation in 100% ethanol (Merck Millipore) at −20 °C overnight, followed by a second precipitation in 100% ethanol at −80°C for 2-4 h. As a quality check for total RNA extraction, agarose gel electrophoresis (1% agarose gel, run 30 min at 100 V) was performed to check for the presence of the 25s, 18s and 5.8s ribosomal RNA bands. Concentration and purity of the RNA samples were measured with a spectrophotometer (Thermo Fisher Scientific, NanoDrop 2000) at 260 nm and 280 nm. cDNA was then obtained from 800 ng total RNA in a PCR cycler (Applied Biosystems, ProFlex PCR system 3×32-well) using random primers and a high-capacity cDNA reverse transcription kit following the included protocol (Thermo Fisher Scientific).

Quantitative PCR (qPCR) measurements were carried out on a LightCycler 480 Multiwell Plate 96 (Roche) using a DNA-binding fluorescent dye (BioRad, SsoAdvanced Universal SYBR Green Supermix) and mRNA sequence specific primers (Sigma-Aldrich). The qPCR was performed with 2 μL of a 1:10 dilution of the cDNA for the genes *ACT1, HHO1, HTB2* and *mCitrine*, or a 1:100 dilution for all other genes. Melting curve data were analyzed to verify primer specificity. Each sample was measured in technical duplicates and the mean value 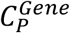 was used for further analyses if 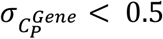. Relative concentrations, normalized on the reference gene *RDN18* were calculated using the equation:

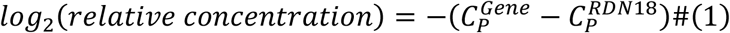

In order to analyze relative concentrations as a function of cell volume, the mean cell volumes were determined from the measured cell volume distributions.

### Test for qPCR primer specificity

To test the specificity of the qPCR primer used to quantify histone mRNA concentrations, we analyzed deletion strains, where possible, for their respective deleted gene to check for unspecific primer binding. For example, we performed a qPCR measurement with the *HHO1* primers on a *hho1Δ* strain and compared the obtained *C_p_* values with the *C_p_* values obtained in the reference strain MS63-1 (**Supplementary Table 1**). We constructed deletion strains for the genes *HHO1, HTB2, HHF1, HHF2, HHT1* and *HHT2*, for which we obtained viable colonies without dramatic growth defects. RNA was extracted as described above, and 1 μg of total RNA was reverse-transcribed using the above mentioned high capacity cDNA synthesis kit. The qPCR was performed with 2 μL of a 1:10 dilution of each cDNA sample, and measured in 3 or 6 technical replicates. *C_p_* values and melting curve data were analyzed to verify primer specificity. Results are shown in **Supplementary Table 2**, deletion strains used are listed in **Supplementary Table 1**, a list of all qPCR primers used can be found in **Supplementary Table 3**.

### Flow Cytometry

Cultures (2 mL – 5 mL) were grown in YPD for at least 6 h in a shaking incubator (30°C, 250 rpm) before being washed and transferred to SCGE and grown for at least 16 h. Appropriate β-estradiol concentrations were then added to inducible cells (0 nM and 30 nM for haploids or 50 nM for diploids), and the cultures grown for at least 24 h in a final volume of 3 mL – 5 mL. During cell growth, *OD*_600_ < 1.3 was maintained through appropriate dilutions.

Cell volume distributions of cultures were measured with a Coulter counter after sonication for 5 s. Cells were fixed using a 37% formaldehyde solution (Sigma-Aldrich) by pipetting 100 μL of formaldehyde into 900 μL of cell cultures in order to achieve a final formaldehyde concentration of 3.7%. Cultures were incubated at room temperature on a rotator (VWR International, Tube Rotator) for 15 min, spun down at 10k g-force for 3 min and subsequently washed and resuspended in 100 μL - 1000 μL 100mM potassium phosphate (pH 7.5). Samples were then stored on ice until being used for flow cytometry.

Flow Cytometry measurements were carried out on a benchtop flow cytometer with octagon and trigon detector arrays (BD Biosciences, LSR II). Strains expressing the fluorescent protein *mCitrine* were excited with a 488 nm coherent sapphire solid-state laser paired with a 530/30 nm filter set. Side-scatter voltage was set to 220 V for all measurements, voltages for forward-scatter and photomultiplier tubes were adjusted depending on whether haploid or diploid cells or both were being measured. However, identical settings were used for replicate experiments. After removing obvious outliers or potential doublets through standard gating strategies, at least 10.000 cells were imaged in the final stopping gate. For each experiment, cells not expressing *mCitrine* were measured to determine the cell-volume-dependent autofluorescence background which was subtracted from the mean fluorescence intensity of each sample measured in the same experiment. In order to calculate fluorescence concentrations, mean cell volumes were determined from the cell volume distributions measured with the Coulter counter. Mean fluorescence concentrations were then calculated by dividing the mean fluorescence intensity of each sample by its mean cell volume, allowing us to analyze mCitrine fluorescence concentrations as a function of cell volume.

### Cell cycle analysis using flow cytometry

To get insights into the distributions of cell cycle phases in cell populations of non-inducible and inducible *WHI5* haploid and diploid strains, we performed cell cycle analysis using flow cytometry. For this purpose, cell cultures (5 mL) were grown in YPD for at least 6 h in a shaking incubator (30*°C*, 250 rpm), before being washed and transferred to SCGE; where appropriate β-estradiol concentrations were added (10 nM or 30 nM for haploid cells, 50 nM for diploid cells). The cultures were then grown for at least 24 h, assuring *OD*_600_ < 1.3 during culture growth through appropriate dilutions. Cell volume distributions of cultures were measured with a Coulter counter after sonication for 5 s. To fixate the cells and subsequently stain the DNA, we followed an already established protocol^41^. Specifically, 1 mL of each cell culture was pipetted into 9 mL of cold 80% ethanol and incubated at 4°C on a rotator overnight. The cultures were then spun down at 4000 rpm for 2 min and washed twice in 50 mM Tris-HCl (pH = 8.0). Cells were then successively treated with a 1 mg/mL RNase A (Thermo Fisher Scientific) solution for 40 min at 37°C, a 20 mg/mL Proteinase K (Promega) solution for 1 h at 37°C and a 10x SYBR Green I (Sigma-Aldrich) solution for 1 h at room temperature. Between each treatment, cells were washed twice with 50 mM Tris-HCl and resuspended in 50 mM Tris-HCl. After the last treatment, cells were sonicated for 5 s. Flow Cytometry measurements were carried out on the benchtop flow cytometer described above, using the same laser, filter sets and side-scatter voltage. Settings for forward-scatter and photomultiplier tubes were adjusted depending on the condition measured. To estimate cell-cycle fractions, imaged DNA content frequency histograms were analyzed using Watson modelling. However, we noticed that for cell populations with large cell volumes (*i.e*. high β-estradiol concentrations), the DNA content distributions showed pronounced tails at large cell volumes that were not fit by the model. We speculate that this tail represents an increased mitochondrial DNA content in large cells^42^, which suggests that a fraction of G1 cells would be wrongly identified as S phase. Thus, we decided to limit our analysis to classifying cells as either G1/S-phase or G2/M-phase (**Supplementary Fig. 1c**). Using this approach, we did not find a drastic influence of the β-estradiol concentration used for Whi5 induction on the cell cycle distributions.

### Volume-dependence parameter

Analyzing protein and mRNA concentrations as a function of cell volume reveals a decrease of concentration with increasing cell volume for histones. In order to quantify this decrease, we performed a linear regression on the double logarithmic data and define the slope of the fit as the *volume-dependence parameter* (VDP):

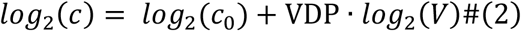

The VDP gives us a quantitative measure for the relation of protein and mRNA concentrations with cell volume: A negative VDP indicates a decrease of concentration with increasing cell volume. The special case of VDP = −1 corresponds to a decrease of concentration with *c*~1/*V*, and therefore signifies a constant amount of protein or mRNA with increasing cell volume. A positive VDP indicates an increase of concentration with increasing cell volume, and VDP = 0 corresponds to a constant concentration *c*_0_.

### Statistical analyses

#### Significance of VDPs

To test for a significant deviation of the VDP from 0, we performed two-tailed one-sample t-tests on the regression coefficients of the linear fit at a confidence level of *α* = 0.05. Our null hypothesis *H*_0_ assumes the respective coefficient to be equal to 0. In order to test for the significance of the VDP, we are interested in the slope of the linear fit: for a p-value smaller than *α*, we reject *H*_0_ and consider the slope, *i.e*. the VDP, to be significantly different from 0.

To test whether the VDPs of two different conditions significantly deviate from each other, we used a general linear regression model with a categorical variable, *Type*, to differentiate between the two conditions analysed:

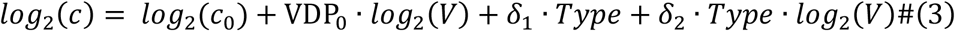

with *c*_0_ and VDP_0_ corresponding to the reference condition (*Type* = 0), *δ*_1_ describing the average difference in the intercepts of the linear fits between the two conditions, and *δ*_2_ describing the change in the slopes (VDPs) between the two conditions. In order to test for a significant difference between the two VDPs, we perform a two-tailed one-sample t-test on *δ*_2_, with the null hypothesis *H*_0_ assuming *δ*_2_ = 0, at a confidence level of *α* = 0.05. For a p-value smaller than *α*, we reject *H*_0_ and consider the change between the two slopes to be significant, *i.e*. we consider the two VDPs to be significantly different from each other.

#### Error estimation of concentrations at 60 fL

To calculate concentrations at a characteristic cell volume of 60 fL with respective error estimates, we evaluated the linear fits to the double logarithmic data at 60 fL and estimated the 95 % confidence intervals of the fit at 60 fL. When normalizing the concentration to a chosen value *x*, errors were calculated using error propagation:

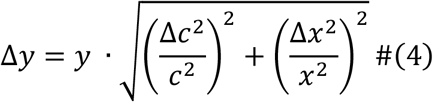

with *y* being the new normalized concentration and *c* the previously calculated concentration.

To estimate the error associated with the ratio between the concentrations at 60 fL in haploids and diploids, we used bootstrap analysis. Specifically, we treated the measurements of protein or mRNA concentration and corresponding cell volume as a set of linked variables, both for haploid and diploid cells. We then resampled n = 10000 populations of same size by random sampling with replacement from this experimental two-dimensional population. Next, we performed a linear regression on the double logarithmic data for each of the resampled populations and estimated the concentration at 60 fL, giving us a distribution of n = 10000 concentrations at 60 fL for both haploid and diploid cells. Finally, we randomly selected a concentration in each of those distributions, and divide the concentration for diploids by the concentration for haploids. We repeated this process 10000 times with replacement to obtain a distribution of n = 10000 concentration ratios, for which we calculate the median and the 2.5- and 97.5-percentiles.

### Minimal model

To obtain mechanistic insight on how the transcription rate of one specific promoter depends on cell volume and ploidy context, we sought to build a minimal model. For this, we consider transcription being limited by one component of the transcriptional machinery, potentially a subunit of the RNA polymerase. In addition, we assume transcript degradation to be the same for all transcripts, and set the corresponding degradation rate *k_deg_* = 1, *i.e*., all other rates are normalized with respect to *k_deg_*. Note that in the case of stable transcripts, *k_deg_* also describes dilution of transcripts by cell growth.

To account for the competition of different promoters for a finite number of the limiting component of the transcriptional machinery (*TM*), our model distinguishes two classes of promoters - a *general pool* of promoters, *g*, with *n_h_* ≈ 6000 (haploids) or *n_d_* ≈ 12000 (diploids), and a single *promoter of interest, p*, present as a single copy. We describe each promoter as one single binding site for *TM* and denote the number of *TM* bound to general promoters as *R^g^*. Binding of *TM* at the single promoter of interest is described by *R^p^*, which can assume values between 0 (not bound) and 1 (bound). Moreover, *R_f_* denotes the number of free *TM*. We assume that the total number of *TM* (free and bound) scales proportionally to cell volume *V* and is given by

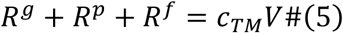

with *c_TM_* being the total *TM* concentration.

Assuming that the arrival of *TM* at promoters is proportional to the concentration of free *TM*, the change in number of bound general promoters over time is given by following equation:

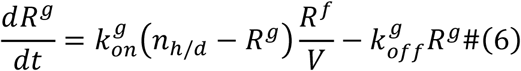

where 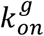 is the rate at which transcription is being initiated at each general promoter, *n_h/d_ – R^g^* are the number of general promoters not bound to *TM* in haploids or diploids, respectively, and 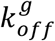 models the rate at which bound *TM* complete transcriptional elongation.

Similarly, the change in binding of *TM* to the single promoter of interest over time is given by:

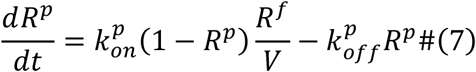

with parameters 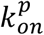 and 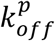 representing transcriptional initiation and elongation, respectively, at the promoter of interest.

Solving (6) and (7) at steady-state 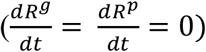, constraints the number of bound *TM*s via the following nonlinear equations

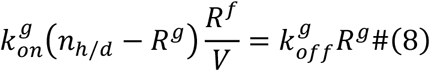

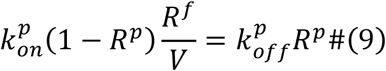

Finally, the steady-state concentration of transcripts produced from the single promoter of interest is equal to 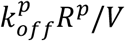.

Given a set of parameters 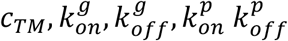, numerically solving equations (5), (8) and (9) allows to calculate the transcript concentration, generated by the single promoter of interest as a function of cell volume *V*. We set *c_TM_* = 2000, 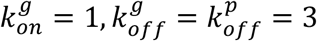 and calculate the steady-state concentration in haploids and diploids over cell volume for 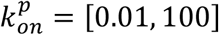.

In order to determine the VDP as a function of 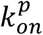, we calculated the concentration for each value of 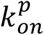 over a cell volume range of 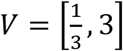 and performed a linear regression fit on the logarithm of the concentration as a function of the logarithm of the cell volume, with cell volumes being equally spaced on the log scale. The VDP is then determined as the slope of the linear fit.

## Data availability statement

Yeast strains and raw data are available upon reasonable request.

## Code availability statement

Additional information on image analysis approaches described in the methods and previous publications is available upon reasonable request.

## Acknowledgments

We thank Matthew Swaffer and Anika Seel for sharing strains, Thomas Hofer and Elfriede Nößner from the HMGU-Immunoanalytics-Core Facility for support with flow cytometry, and the Institute of Functional Epigenetics and the Skotheim lab for discussions. We thank Matthew Swaffer, Amanda Amodeo and Jan Skotheim for comments on the manuscript. This work was supported by the DFG through project SCHM3031/4-1, by the Human Frontier Science Program (career development award to K.M.S) as well as the AMPro program (ZT0026) and the Helmholtz Gesellschaft. A.S. was supported by NIH grants 5R01GM124446 and 5R01GM126557.

## References

1. Marguerat, S. & Bähler, J. Coordinating genome expression with cell size. Trends Genet. 28, 560–565 (2012).

2. Schmoller, K. M. & Skotheim, J. M. The Biosynthetic Basis of Cell Size Control. Trends Cell Biol. 25, 793–802 (2015).

3. Zhurinsky, J. et al. A coordinated global control over cellular transcription. Curr. Biol. 20, 2010–2015 (2010).

4. Padovan-Merhar, O. et al. Single Mammalian Cells Compensate for Differences in Cellular Volume and DNA Copy Number through Independent Global Transcriptional Mechanisms. Mol. Cell 58, 339–352 (2015).

5. Sun, X.-M. et al. Size-dependent increase in RNA Polymerase II initiation rates mediates gene expression scaling with cell size. bioRxiv Mol. Biol. (2019) doi:10.1101/754788.

6. Nadal-Ribelles, M. et al. Sensitive high-throughput single-cell RNA-seq reveals within-clonal transcript correlations in yeast populations. Nat. Microbiol. (2019) doi:10.1038/s41564-018-0346-9.

7. Amodeo, A. A., Jukam, D., Straight, A. F. & Skotheim, J. M. Histone titration against the genome sets the DNA-to-cytoplasm threshold for the Xenopus midblastula transition. Proc. Natl. Acad. Sci. U. S. A. (2015) doi:10.1073/pnas.1413990112.

8. Joseph, S. R. et al. Competition between histone and transcription factor binding regulates the onset of transcription in zebrafish embryos. Elife (2017) doi:10.7554/eLife.23326.

9. Hauer, M. H. et al. Histone degradation in response to DNA damage enhances chromatin dynamics and recombination rates. Nat. Struct. Mol. Biol. (2017) doi:10.1038/nsmb.3347.

10. Chari, S., Wilky, H., Govindan, J. & Amodeo, A. A. Histone concentration regulates the cell cycle and transcription in early development. Dev. (2019) doi:10.1242/dev.177402.

11. Kim, U. J., Han, M., Kayne, P. & Grunstein, M. Effects of histone H4 depletion on the cell cycle and transcription of Saccharomyces cerevisiae. EMBO J. (1988) doi:10.1002/j.1460-2075.1988.tb03060.x.

12. Han, M., Chang, M., Kim, U. J. & Grunstein, M. Histone H2B repression causes cell-cycle-specific arrest in yeast: Effects on chromosomal segregation, replication, and transcription. Cell (1987) doi:10.1016/0092-8674(87)90237-6.

13. Meeks-Wagner, D. & Hartwell, L. H. Normal stoichiometry of histone dimer sets is necessary for high fidelity of mitotic chromosome transmission. Cell 44, 43–52 (1986).

14. Eriksson, P. R., Ganguli, D., Nagarajavel, V. & Clark, D. J. Regulation of histone gene expression in budding yeast. Genetics 191, 7–20 (2012).

15. Kurat, C. F. et al. Regulation of histone gene transcription in yeast. Cell. Mol. Life Sci. 71, 599–613 (2014).

16. Gunjan, A. & Verreault, A. A Rad53 Kinase-Dependent Surveillance Mechanism that Regulates Histone Protein Levels in S. cerevisiae. Cell 115, 537–549 (2003).

17. Doncic, A., Eser, U., Atay, O. & Skotheim, J. M. An Algorithm to Automate Yeast Segmentation and Tracking. PLoS One 8, (2013).

18. Schmoller, K. M., Turner, J. J., Kõivomägi, M. & Skotheim, J. M. Dilution of the cell cycle inhibitor Whi5 controls budding-yeast cell size. Nature 526, 268–272 (2015).

19. Ottoz, D. S. M., Rudolf, F. & Stelling, J. Inducible, tightly regulated and growth condition-independent transcription factor in Saccharomyces cerevisiae. Nucleic Acids Res. 42, (2014).

20. Norris, D. & Osley, M. A. The two gene pairs encoding H2A and H2B play different roles in the Saccharomyces cerevisiae life cycle. Mol. Cell. Biol. (1987) doi:10.1128/mcb.7.10.3473.

21. Moran, L., Norris, D. & Osley, M. A. A yeast H2A-H2B promoter can be regulated by changes in histone gene copy number. Genes Dev. (1990) doi:10.1101/gad.4.5.752.

22. Cross, S. L. & Smith, M. M. Comparison of the structure and cell cycle expression of mRNAs encoded by two histone H3-H4 loci in Saccharomyces cerevisiae. Mol. Cell. Biol. 8, 945–54 (1988).

23. von der Haar, T. A quantitative estimation of the global translational activity in logarithmically growing yeast cells. BMC Syst. Biol. (2008) doi:10.1186/1752-0509-2-87.

24. Williamson, D. H. & Scopes, A. W. The distribution of nucleic acids and protein between different sized yeast cells. Exp. Cell Res. (1961) doi:10.1016/0014-4827(61)90258-0.

25. Mena, A. et al. Asymmetric cell division requires specific mechanisms for adjusting global transcription. Nucleic Acids Res. 45, 12401–12412 (2017).

26. Hereford, L., Fahrner, K., Woolford, J., Rosbash, M. & Kaback, D. B. Isolation of yeast histone genes H2A and H2B. Cell (1979) doi:10.1016/0092-8674(79)90237-X.

27. Smith, M. M. & Murray, K. Yeast H3 and H4 histone messenger RNAs are transcribed from two non-allelic gene sets. J. Mol. Biol. (1983) doi:10.1016/S0022-2836(83)80163-6.

28. Libuda, D. E. & Winston, F. Alterations in DNA replication and histone levels promote histone gene amplification in Saccharomyces cerevisiae. Genetics (2010) doi:10.1534/genetics.109.113662.

29. Fillingham, J. et al. Two-Color Cell Array Screen Reveals Interdependent Roles for Histone Chaperones and a Chromatin Boundary Regulator in Histone Gene Repression. Mol. Cell 35, 340–351 (2009).

30. Zunder, R. M. & Rine, J. Direct Interplay among Histones, Histone Chaperones, and a Chromatin Boundary Protein in the Control of Histone Gene Expression. Mol. Cell. Biol. 32, 4337–4349 (2012).

31. Feser, J. et al. Elevated Histone Expression Promotes Life Span Extension. Mol. Cell (2010) doi:10.1016/j.molcel.2010.08.015.

32. Canavan, R. & Bond, U. Deletion of the nuclear exosome component RRP6 leads to continued accumulation of the histone mRNA HTB1 in S-phase of the cell cycle in Saccharomyces cerevisiae. Nucleic Acids Res. (2007) doi:10.1093/nar/gkm691.

33. Beggs, S., James, T. C. & Bond, U. The PolyA tail length of yeast histone mRNAs varies during the cell cycle and is influenced by Sen1p and Rrp6p. Nucleic Acids Res. 40, 2700–2711 (2012).

34. Wu, C. Y., Alexander Rolfe, P., Gifford, D. K. & Fink, G. R. Control of transcription by cell size. PLoS Biol. 8, (2010).

35. Osley, M. A., Gould, J., Kim, S., Kane, M. & Hereford, L. Identification of sequences in a yeast histone promoter involved in periodic transcription. Cell 45, 537–544 (1986).

36. Zatulovskiy, E., Berenson, D. F., Topacio, B. R. & Skotheim, J. M. overexpression increased cell size in tissue culture and a mouse cancer model, while. (2018).

37. Chen, Y., Zhao, G., Zahumensky, J., Honey, S. & Futcher, B. Differential Scaling of Gene Expression with Cell Size May Explain Size Control in Budding Yeast. Mol. Cell (2020) doi:10.1016/j.molcel.2020.03.012.

38. Kukhtevich, I. V., Lohrberg, N., Padovani, F., Schneider, R. & Schmoller, K. M. Cell size sets the diameter of the budding yeast contractile ring. Nat. Commun. (2020) doi:10.1038/s41467-020-16764-x.

39. Chandler-Brown, D., Schmoller, K. M., Winetraub, Y. & Skotheim, J. M. The Adder Phenomenon Emerges from Independent Control of Pre-and Post-Start Phases of the Budding Yeast Cell Cycle. Curr. Biol. 27, 2774–2783.e3 (2017).

40. Collart, M. A. & Oliviero, S. Preparation of Yeast RNA. Curr. Protoc. Mol. Biol. (1993) doi:10.1002/0471142727.mb1312s23.

41. Örd, M., Venta, R., Möll, K., Valk, E. & Loog, M. Cyclin-Specific Docking Mechanisms Reveal the Complexity of M-CDK Function in the Cell Cycle. Mol. Cell (2019) doi:10.1016/j.molcel.2019.04.026.

42. Rafelski, S. M. et al. Mitochondrial network size scaling in budding yeast. Science (80-.). (2012) doi:10.1126/science.1225720.

