## Supplementary Information for "Transcription coordinates histone amounts and genome content"

### Extended Data figures

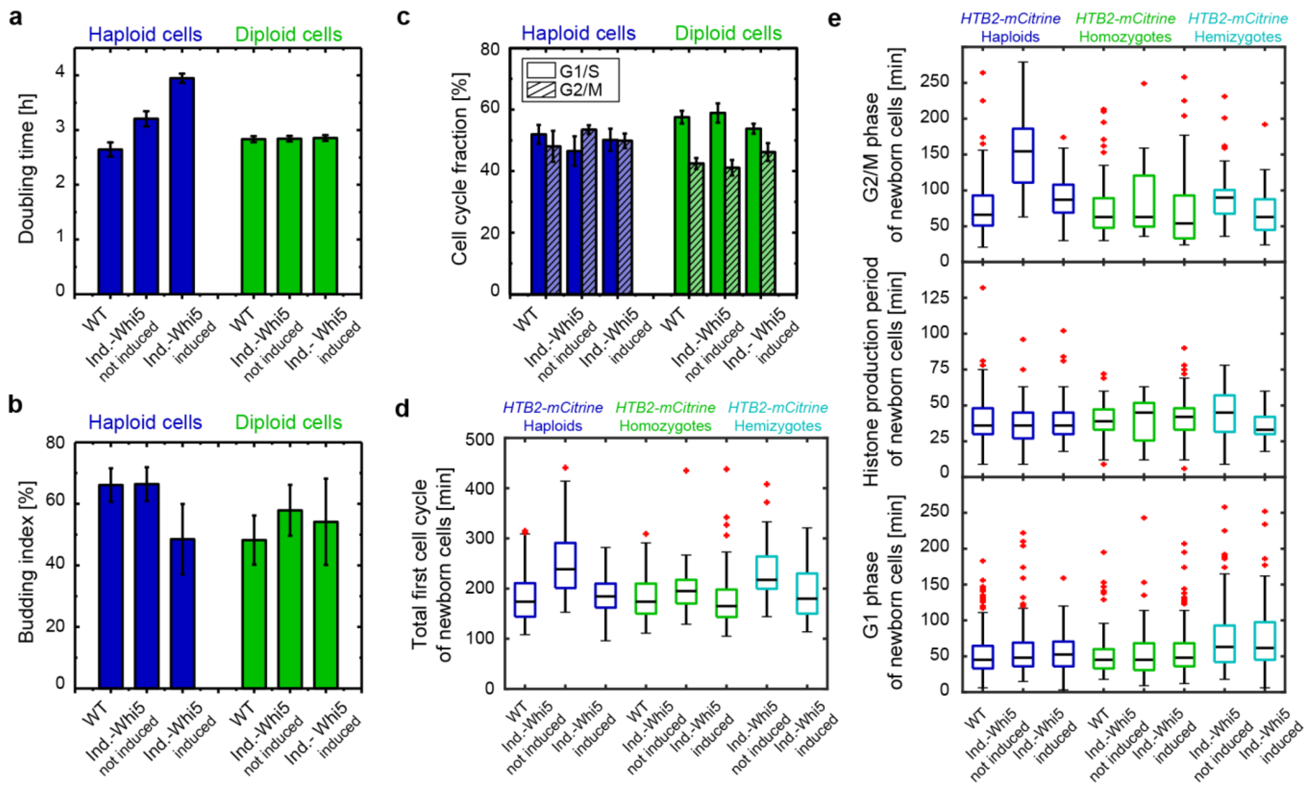


**Supplementary Figure 1.** Overexpression of Whi5 does not drastically change doubling times, budding indices or cell cycle distributions. In all panels, haploid cells were induced with 30 nM β-estradiol, diploid cells with 50 nM. (a) Doubling times calculated from growth curves of exponentially growing cell populations for inducible and non-inducible (WT), untagged haploids (blue) and diploids (green). Error bars indicate the standard deviation of n = 3 biological replicates, n = 2 for induced haploid cells. (b) Budding index (percentage of budded cells), calculated by counting of budded and non-budded cells in exponentially growing cell populations using an optical microscope, for inducible and non-inducible (WT), untagged haploids (blue) and diploids (green). Error bars indicate the standard deviation for n = 4 biological replicates for inducible haploids, n = 3 for the wildtype haploids, and n = 2 for all the diploid strains. (c) Cell cycle distributions (percentage of cells in G1/S-phases, or in G2/M phases) of inducible and non-inducible (WT), untagged haploid (blue) and diploid (green) cells, calculated from population distributions obtained through SYBR Green I staining of the DNA and flow cytometry. Error bars indicate the standard deviation of n = 6 biological replicates. (d & e)

Total first cell cycles of new-born cells (d) as well as individual G1- & G2/M-phases and histone production periods of new-born cells (e) for inducible and non-inducible (WT) *HTB2-mCitrine* haploids (blue), *HTB2-mCitrine* homozygous diploids (green) and *HTB2-mCitrine/htb2Δ* hemizygous diploids calculated from single cell fluorescent traces, measured with live cell fluorescence microscopy. G1 phases are defined as the time from birth to the first budding, histone production phases as the time between the first point of increase in fluorescent signal and the last point of increase, G2/M phases as the time from end of histone production phase to the separation of the cell from its bud. Black, horizontal lines indicate the median between single cells ($n_{haploid}^{WT} = 145, n_{haploid}^{not\ ind.} = 58, n_{haploid}^{ind.} = 58, n_{homoz}^{WT} = 72, n_{homoz.}^{not\ ind.} = 21, n_{homoz.}^{ind} = 85, n_{hemiz.}^{not\ ind.} = 47$and $n_{hemiz.}^{ind.} = 43$), coloured boxes highlight the 25- and 75-percentiles, whiskers extend to $\pm 2.7\sigma$ of the distribution and red crosses highlight outliers.

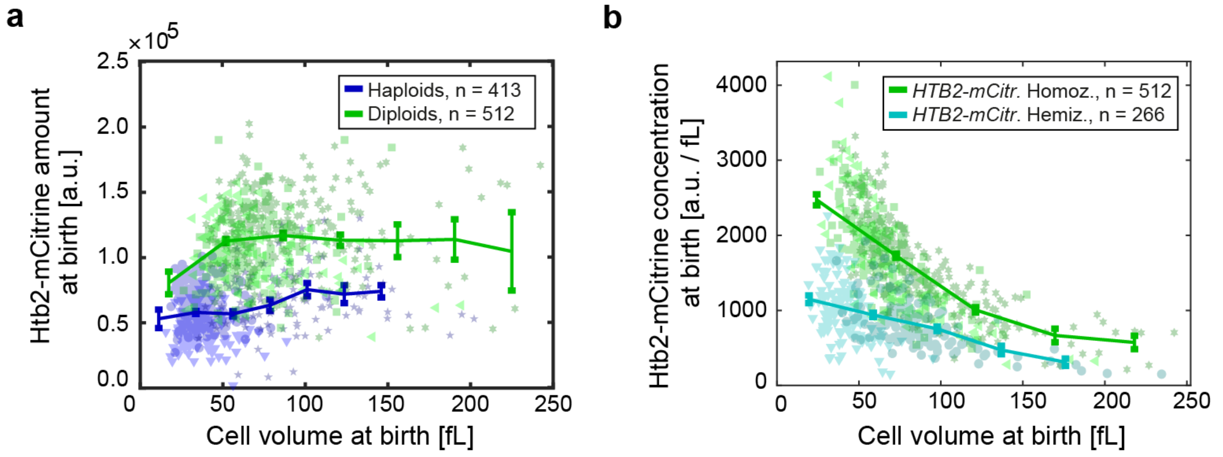


**Supplementary Figure 2.** Htb2-mCitrine amounts and concentrations increase with ploidy. (a) Htb2-mCitrine amounts at birth for *HTB2-mCitrine* haploids and *HTB2-mCitrine* homozygous diploids estimated from fluorescence microscopy as a function of cell volume. Individual data points for the different conditions (▼ 0 nM, ● non-inducible, ★ 30 nM, for haploids and ◀ 0 nM, ■ non-inducible, ✶ 50 nM, for diploids) are highlighted in blue (haploids) and green (diploids). Lines connect binned means, error bars indicate standard errors. (b) Htb2-mCitrine concentrations at birth for *HTB2-mCitrine* homozygous diploids and *HTB2-mCitrine/htb2Δ* hemizygous diploids as a function of cell volume. Individual data points for the different conditions (▼ 0 nM, ● 50 nM, for *HTB2-mCitrine/htb2Δ* (hemizygotes) and ◀ 0 nM, ■ non-inducible, ✶ 50 nM, for *HTB2-mCitrine* homozygotes) are highlighted in teal (hemizygotes) and green (homozygotes). Lines connect binned means, error bars indicate standard errors.

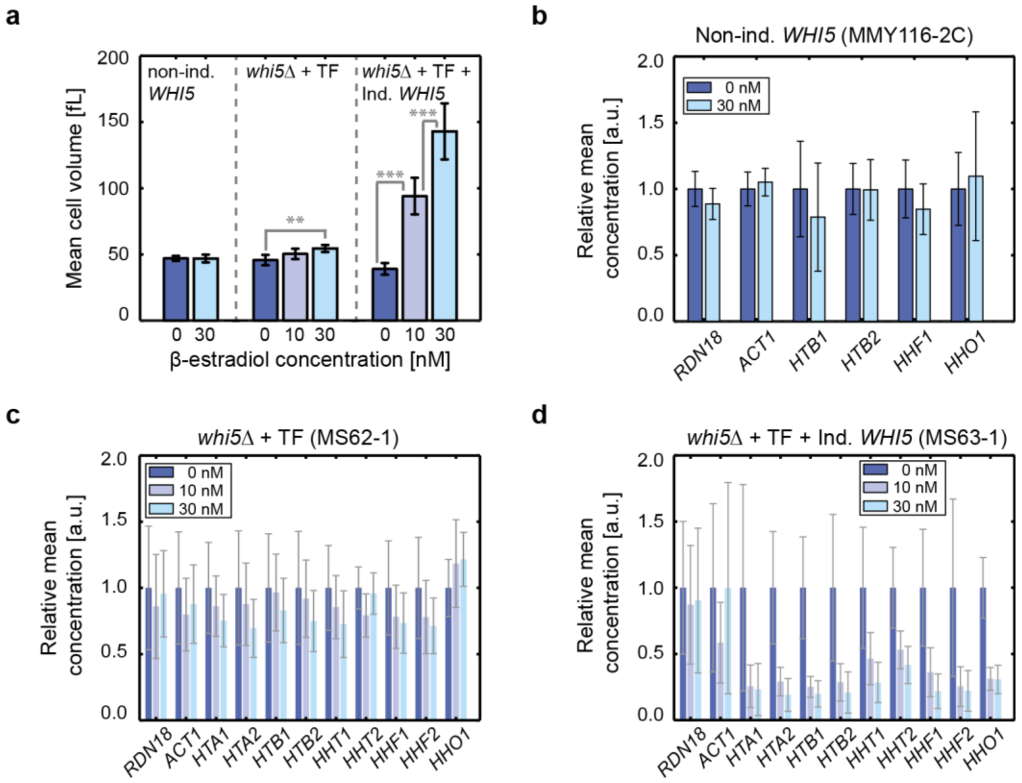


**Supplementary Figure 3.** Decrease in histone transcript concentration with cell volume is specific to Whi5-dependent cell volume increase. (a) Mean cell volumes of exponentially growing cell populations measured with a Coulter counter are shown as a function of β-estradiol concentrations for non-inducible haploid cells, *whi5Δ* haploid cells with β-estradiol-dependent transcription factor (TF), and *whi5Δ* haploid cells with β-estradiol-dependent transcription factor (TF) and β-estradiol-inducible *WHI5*. Error bars indicate the standard deviation of $n^{0}_{non-ind.} = 11$, $n^{30}_{non-ind,} = 4$, $n^{0}_{whi5\Delta+TF} = 5$, $n^{10}_{whi5\Delta+TF} = 6$, $n^{30}_{whi5\Delta+TF} = 5$, $n^{0}_{Ind.-WHI5} = 11$, $n^{10}_{Ind.-WHI5} = 9$, $n^{30}_{Ind.-WHI5} = 10$ biological replicates. Significances were tested using two-tailed two-sample t-tests, after checking for normal distribution and equal variance distributions using a Shapiro-Wilk test and a Bartlett test, respectively, **p<0.01, ***p<0.001. (b-d) Relative mean mRNA concentration of exponentially growing haploid cell populations with and without β-estradiol addition. (b) Non-inducible cells, (c) *whi5Δ* cells with β-estradiol-dependent transcription factor (TF), (d) *whi5Δ* cells with β-estradiol-dependent transcription factor (TF) and β-estradiol inducible *WHI5*. For each gene, values are normalized on the mean mRNA concentration of the cell populations without β-estradiol addition (0 nM). Error bars are derived by error propagation of the standard deviation of $n = 4$ (b), $n \geq 5$ (c), and $n \geq 9$ (d) biological replicates.

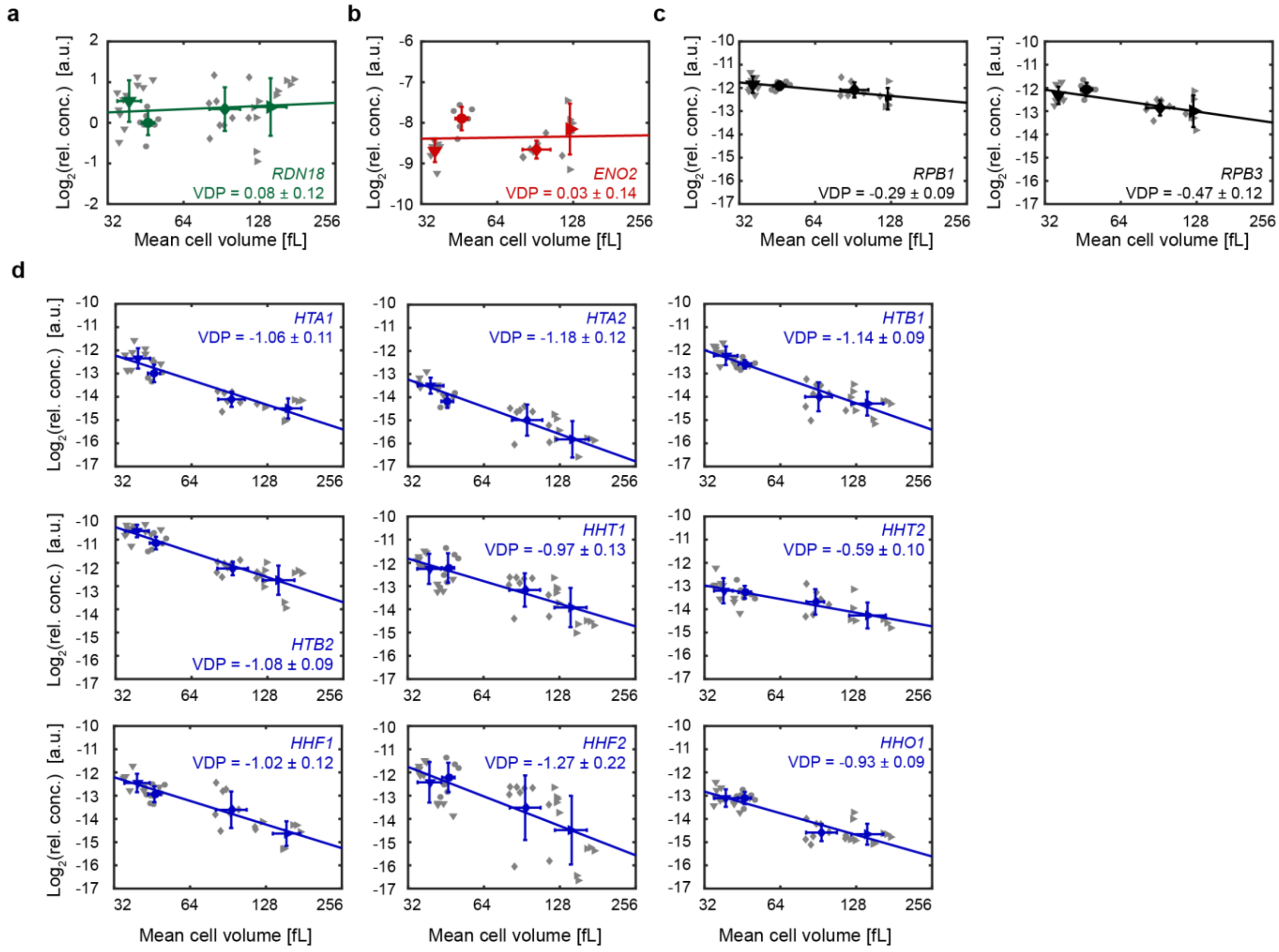


**Supplementary Figure 4.** Histone mRNA concentrations measured by RT-qPCR decrease with increasing cell volume. (a) Relative *RDN18* mRNA concentration for non-inducible and inducible haploid cells over mean cell volume, shown in a double logarithmic plot. mRNA concentrations are normalized on the mean mRNA concentration of non-inducible cells (●). (b - d) Raw data corresponding to Fig. 2d: Relative mRNA concentrations of *ENO2* (b), *RPB1* and *RPB3* (b), as well as all core histone genes and *HHO1* (d) for non-inducible and inducible haploid cells over mean cell volume, shown in double logarithmic plots. mRNA concentrations are normalized on *RDN18* concentrations. Individual data points for the different conditions (▼ 0 nM, ● non-inducible, ◆ 10 nM, ► 30 nM) are highlighted in grey. Green (b), red (c), black (d) or blue (e) symbols indicate the mean of the different conditions. Error bars indicate standard deviations for $n \geq 5$ biological replicates.

Lines show linear fits to the double logarithmic data, with volume-dependence parameters (VDPs) determined as the slope of the fit, with respective standard error.

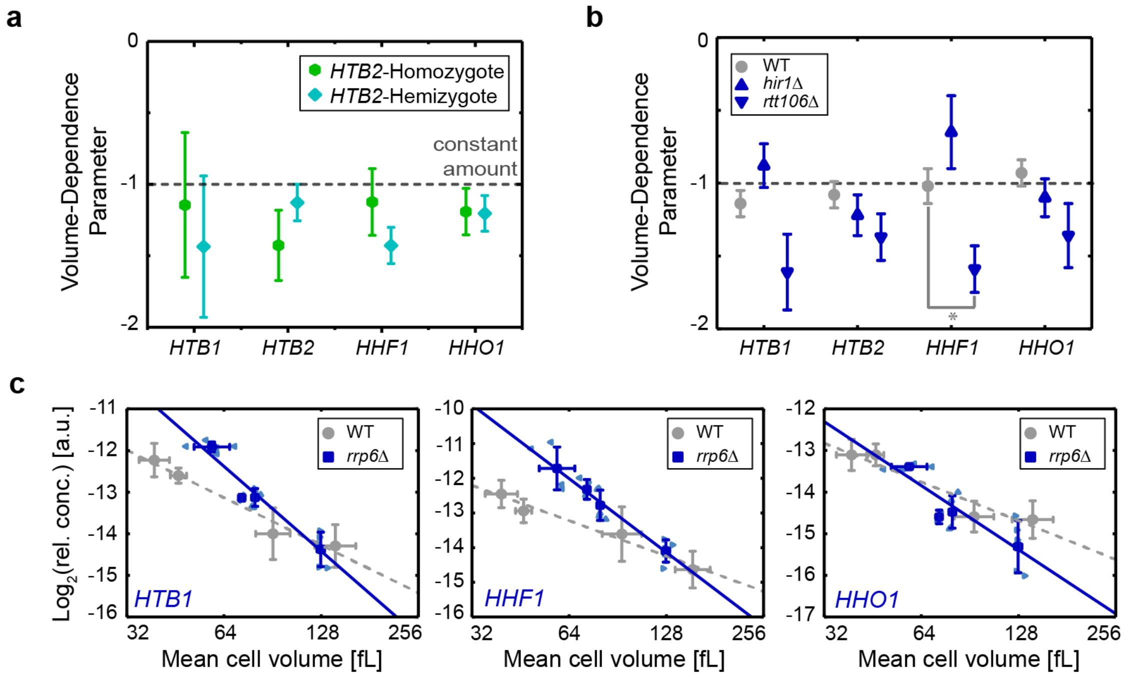


**Supplementary Figure 5.** Direct feedback mechanisms, Hir1-dependent feedback, as well as 3'- to 5'-end degradation by nuclear exosome is not necessary for the decrease of histone mRNA concentrations. (a & b) Summaries of the VDPs determined by RT-qPCR for *HTB1*, *HTB2*, *HHF1* and *HHO1*. VDPs were determined as the slopes of the linear fits to the double logarithmic concentration over cell volume data. Error bars indicate the standard error of the slope. (a) *HTB2* homozygous (green ●) and *HTB2/htb2*Δ hemizygous (teal ◆). (b) Cells carrying no deletion (WT, grey ●), *hir1*Δ (blue ▲) and *rtt106*Δ (blue ▼) cells. Significant VDP variation from the WT were determined using linear regressions, *$p<0.05$. (c) Raw data corresponding to Fig. 2g: Relative mRNA concentrations (normalized on *RDN18*) for inducible and non-inducible haploid cells over mean cell volume, shown in double logarithmic plots. Data corresponding to the *rrp6*Δ cells are highlighted in blue. Light blue symbols highlight the different conditions (◆ non-inducible, ◀ 0 nM, ▲ 10 nM, ▶ 30 nM). Dark blue symbols (■) indicate the mean for each condition. Grey symbols (●) indicate the mean for each condition of the wildtype (carrying no deletion). Lines show the linear fits to the double logarithmic data.

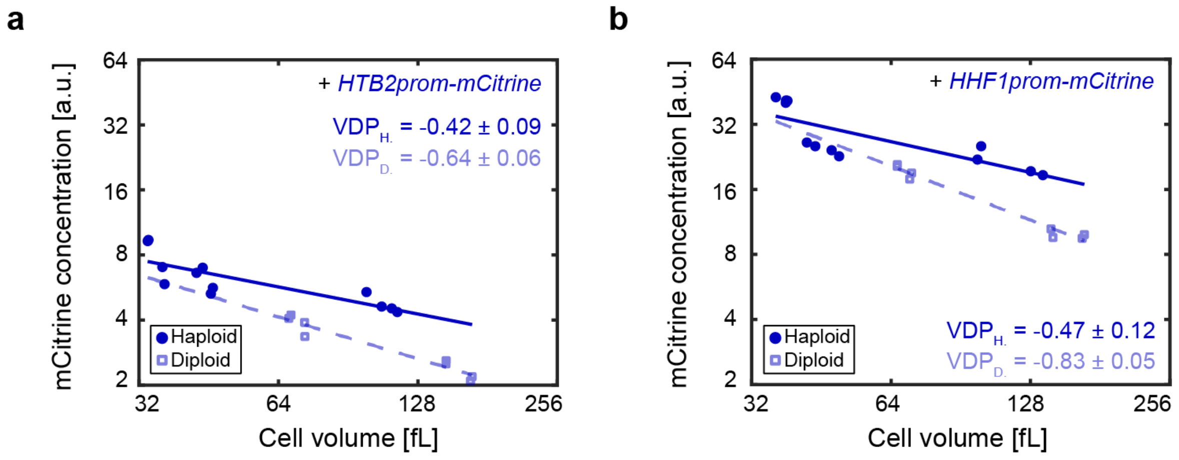


**Supplementary Figure 6** Concentration of mCitrine expressed from a single additional histone promoter in haploid and diploid cells measured with flow cytometry. *mCitrine* concentration, driven by an additional copy of the *HTB2* (a) or *HHF1* (b) promoter in haploid (blue ●) and diploid (blue □) cells, shown as a function of cell volume in a double logarithmic plot. Lines show linear fits to the double logarithmic data with volume-dependence parameters (VDPs) determined as the slope of the fit, with respective standard error.

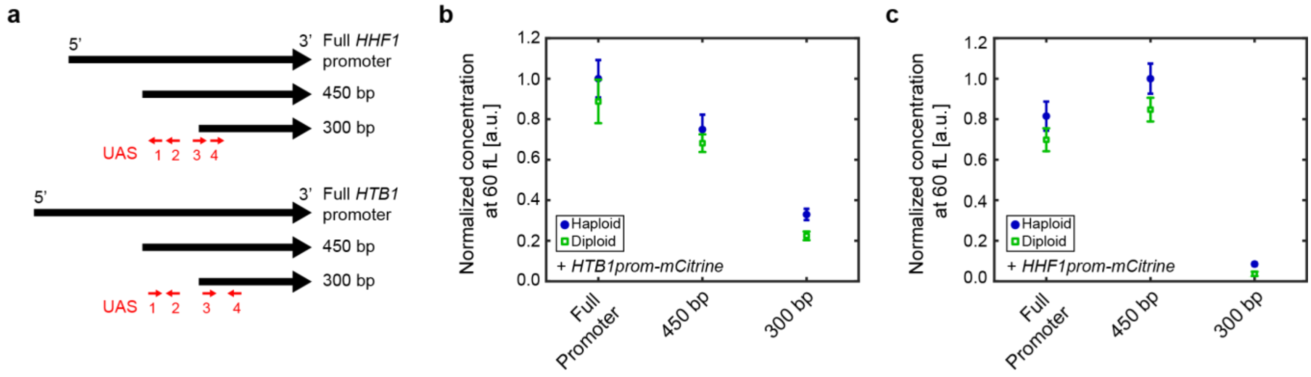


**Supplementary Figure 7.** mCitrine expression driven by single additional histone promoter truncations and measured by flow cytometry decreases once part of the upstream activating sequences (UASs) are truncated. (a) Illustration of the full *HHF1* and *HTB1* promoter, as well as the 450 bp and 300 bp truncations. Red arrows show the location of the upstream activating sequences (UASs), known from the literature[35]. (b - c) Normalized mCitrine concentration at 60 fL for different histone promoter truncations, integrated in haploid (blue ●) and diploid (green □) cells. (b) *mCitrine* driven by *HTB1* promoter truncations, (c) *mCitrine* driven by *HHF1* promoter truncations. Concentrations were calculated by the linear fit to the double logarithmic concentration over cell volume data at 60 fL, and normalized to the maximum concentration calculated for haploid cells. Error bars are derived by error propagation of the 95% confidence interval of the linear fit at 60 fL.

#### Extended Data tables

| **Name** | **Genotype** | **Description** | **Origin** | **Fig.** |
|---|---|---|---|---|
| ASY020-1 | *Mat α/a; ADE2/ADE2, URA3/ura3, leu2/LEU2* | Non-inducible *WHI5*, diploid strain | Anika Seel, Schmoller lab | 1, S1 |
| DBY002-1 | *Mat a; ADE2, whi5Δ::kanMX6-LexAprom-WHI5-ADH1term-LEU2, his3::LexA-ER-AD-TF-HIS3, htb2Δ::KlacURA3* | Inducible *WHI5*, haploid *htb2Δ* strain | This study | Table S2 |
| DBY003-1 | *Mat a; ADE2, whi5Δ::kanMX6-LexAprom-WHI5-ADH1term-LEU2, his3::LexA-ER-AD-TF-HIS3, hho1Δ::CglaTRP1* | Inducible *WHI5*, haploid *hho1Δ* strain | This study | Table S2 |
| DBY008-1 | *Mat a; ADE2, whi5Δ::kanMX6-LexAprom-WHI5-ADH1term-LEU2, his3::LexA-ER-AD-TF-HIS3, hhf1Δ::CglaTRP1* | Inducible *WHI5*, haploid *hhf1Δ* strain | This study | Table S2 |
| DBY009-1 | *Mat α; ADE2, hhf2Δ::CglaTRP1* | Non-inducible *WHI5*, haploid *hhf2Δ* strain | This study | Table S2 |
| DBY011-1 | *Mat α; ADE2, hht1Δ::CglaTRP1* | Non-inducible *WHI5*, haploid *hht1Δ* strain | This study | Table S2 |
| DBY013-1 | *Mat α; ADE2, hht2Δ::CglaTRP1* | Non-inducible *WHI5*, haploid *hht2Δ* strain | This study | Table S2 |
| DBY020-2 | *Mat α; ADE2, ura3::HTB1prom-mCitrine-ADH1term-CglaTRP1-URA3* | Non-inducible *WHI5*, haploid strain with additional copy of *HTB1* promoter expressing *mCitrine* | This study | 3, 5, S6, S7 |
| DBY021-3 | *Mat α; ADE2, ura3::HTB2prom-mCitrine-ADH1term-CglaTRP1-URA3* | Non-inducible *WHI5*, haploid strain with additional copy of *HTB2* promoter expressing *mCitrine* | This study | 3, S6 |
| DBY022-1 | *Mat α; ADE2, ura3::HHF1prom-mCitrine-ADH1term-CglaTRP1-URA3* | Non-inducible *WHI5*, haploid strain with additional copy of *HHF1* promoter expressing *mCitrine* | This study | 3, 5, S6, S7 |
| KCY001-3 | *Mat a; ADE2, htb2::HTB2-linker-mCitrine-ADH1term-CglaTRP1, whi5Δ::kanMX6-LexAprom-WHI5-ADH1term-LEU2, his3::LexA-ER-AD-TF-HIS3* | Inducible *WHI5*, haploid strain with *HTB2* tagged with *mCitrine* | This study | 1, S1, S2 |
| KCY002-3 | *Mat α; ADE2, htb2::HTB2-linker-mCitrine-ADH1term-CglaTRP1* | Non-inducible *WHI5*, haploid strain with *HTB2* tagged with *mCitrine* | This study | 1, S1, S2 |
| KCY005-1 | *Mat α/a; ADE2/ADE2, whi5Δ::CglaTRP1/whi5Δ::kanMX6-LexAprom-WHI5-ADH1term-LEU2, his3/his3Δ::LexA-ER-AD-TF-HIS3* | Inducible *WHI5*, diploid strain | This study | 1, 2, S1, S5 |

| Name | Genotype | Description | Origin | Fig. |
|---|---|---|---|---|
| KCY006-1 | *Mat α/a; ADE2/ADE2, htb2Δ::KlacURA3/HTB2, whi5Δ::CglaTRP1/whi5Δ::kanMX6-LexAprom-WHI5-ADH1term-LEU2, HI3/his3Δ::LexA-ER-AD-TF-HIS3* | Inducible *WHI5*, diploid strain with one *HTB2* allele deleted | This study | 2, S5 |
| KCY007-2 | *Mat α; ADE2, ura3::150bpHHF1prom-mCitrine-ADH1term-CglaTRP1-URA3* | Non-inducible *WHI5*, haploid strain with additional 150 bp of *HHF1* promoter (truncated 5' – 3') expressing *mCitrine* | This study | 5 |
| KCY008-1 | *Mat α; ADE2, ura3::300bpHHF1prom-mCitrine-ADH1term-CglaTRP1-URA3* | Non-inducible *WHI5*, haploid strain with additional 300 bp of *HHF1* promoter (truncated 5' – 3') expressing *mCitrine* | This study | 5, S7 |
| KCY009-1 | *Mat α; ADE2, ura3::450bpHHF1prom-mCitrine-ADH1term-CglaTRP1-URA3* | Non-inducible *WHI5*, haploid strain with additional 450 bp of *HHF1* promoter (truncated 5' – 3') expressing *mCitrine* | This study | 5, S7 |
| KCY010-2 | *Mat α; ADE2, ura3::600bpHHF1prom-mCitrine-ADH1term-CglaTRP1-URA3* | Non-inducible *WHI5*, haploid strain with additional 600 bp of *HHF1* promoter (truncated 5' – 3') expressing *mCitrine* | This study | 5 |
| KCY011-1 | *Mat a; ADE2, ura3::150bpHHF1prom-mCitrine-ADH1term-CglaTRP1-URA3, whi5Δ::kanMX6-LexAprom-WHI5-ADH1term-LEU2, his3::LexA-ER-AD-TF-HIS3* | Inducible *WHI5*, haploid strain with additional 150 bp of *HHF1* promoter (truncated 5' – 3') expressing *mCitrine* | This study | 5 |
| KCY012-1 | *Mat a; ADE2, ura3::300bpHHF1prom-mCitrine-Adh1term-CglaTRP1-URA3, whi5Δ::kanMX6-LexAprom-WHI5-ADH1term-LEU2, his3::LexA-ER-AD-TF-HIS3* | Inducible *WHI5*, haploid strain with additional 300 bp of *HHF1* promoter (truncated 5' – 3') expressing *mCitrine* | This study | 5, S7 |
| KCY013-1 | *Mat a; ADE2, ura3::450bpHHF1prom-mCitrine-ADH1term-CglaTRP1-URA3, whi5Δ::kanMX6-LexAprom-WHI5-ADH1term-LEU2, his3::LexA-ER-AD-TF-HIS3* | Inducible *WHI5*, haploid strain with additional 450 bp of *HHF1* promoter (truncated 5' – 3') expressing *mCitrine* | This study | 5, S7 |
| KCY014-1 | *Mat a; ADE2, ura3::600bpHHF1prom-mCitrine-Adh1term-CglaTRP1-URA3, whi5Δ::kanMX6-LexAprom-WHI5-ADH1term-LEU2, his3::LexA-ER-AD-TF-HIS3* | Inducible *WHI5*, haploid strain with additional 600 bp of *HHF1* promoter | This study | 5 |

| Name | Genotype | Description | Origin | Fig. |
|---|---|---|---|---|
| | | (truncated 5' – 3') expressing *mCitrine* | | |
| KCY015-1 | *Mat a; ADE2, ura3::150bpHTB1prom-mCitrine-ADH1term-CglaTRP1-URA3, whi5Δ::kanMX6-LexAprom-WHI5-ADH1term-LEU2, his3::LexA-ER-AD-TF-HIS3* | Inducible *WHI5*, haploid strain with additional 150 bp of *HTB1* promoter (truncated 5' – 3') expressing *mCitrine* | This study | 5 |
| KCY016-1 | *Mat a; ADE2, ura3::300bpHTB1prom-mCitrine-ADH1term-CglaTRP1-URA3, whi5Δ::kanMX6-LexAprom-WHI5-ADH1term-LEU2, his3::LexA-ER-AD-TF-HIS3* | Inducible *WHI5*, haploid strain with additional 300 bp of *HTB1* promoter (truncated 5' – 3') expressing *mCitrine* | This study | 5, S7 |
| KCY017-1 | *Mat a; ADE2, ura3::450bpHTB1prom-mCitrine-ADH1term-CglaTRP1-URA3, whi5Δ::kanMX6-LexAprom-WHI5-ADH1term-LEU2, his3::LexA-ER-AD-TF-HIS3* | Inducible *WHI5*, haploid strain with additional 450 bp of *HTB1* promoter (truncated 5' – 3') expressing *mCitrine* | This study | 5, S7 |
| KCY018-1 | *Mat a; ADE2, ura3::600bpHTB1prom-mCitrine-ADH1term-CglaTRP1-URA3, whi5Δ::kanMX6-LexAprom-WHI5-ADH1term-LEU2, his3::LexA-ER-AD-TF-HIS3* | Inducible *WHI5*, haploid strain with additional 600 bp of *HTB1* promoter (truncated 5' – 3') expressing *mCitrine* | This study | 5 |
| KCY019-1 | *Mat a; ADE2, ura3::750bpHTB1prom-mCitrine-ADH1term-CglaTRP1-URA3, whi5Δ::kanMX6-LexAprom-WHI5-ADH1term-LEU2, his3::LexA-ER-AD-TF-HIS3* | Inducible *WHI5*, haploid strain with additional 750 bp of *HTB1* promoter (truncated 5' – 3') expressing *mCitrine* | This study | 5 |
| KCY020-1 | *Mat α; ADE2, ura3::150bpHTB1prom-mCitrine-ADH1term-CglaTRP1-URA3* | Non-inducible *WHI5*, haploid strain with additional 150 bp of *HTB1* promoter (truncated 5' – 3') expressing *mCitrine* | This study | 5 |
| KCY021-1 | *Mat α; ADE2, ura3::300bpHTB1prom-mCitrine-ADH1term-CglaTRP1-URA3* | Non-inducible *WHI5*, haploid strain with additional 300 bp of *HTB1* promoter (truncated 5' – 3') expressing *mCitrine* | This study | 5, S7 |
| KCY022-1 | *Mat α; ADE2, ura3::450bpHTB1prom-mCitrine-ADH1term-CglaTRP1-URA3* | Non-inducible *WHI5*, haploid strain with additional 450 bp of *HTB1* promoter (truncated 5' – 3') expressing *mCitrine* | This study | 5, S7 |
| KCY023-4 | *Mat α; ADE2, ura3::600bpHTB1prom-mCitrine-ADHterm-CglaTRP1-URA3* | Non-inducible *WHI5*, haploid strain with | This study | 5 |

| Name | Genotype | Description | Origin | Fig. |
|---|---|---|---|---|
| | | additional 600 bp of *HTB1* promoter (truncated 5' – 3') expressing *mCitrine* | | |
| KCY024-1 | *Mat α; ADE2, ura3::750bpHTB1prom-mCitrine-ADH1term-CglaTRP1-URA3* | Non-inducible *WHI5*, haploid strain with additional 750 bp of *HTB1* promoter (truncated 5' – 3') expressing *mCitrine* | This study | 5 |
| KCY027-4 | *Mat α/a; ADE2/ADE2, htb2::HTB2-linker-mCitrine-ADH1term-KlacURA3/htb2::HTB2-linker-mCitrine-ADH1term-CglaTRP1* | Non-inducible *WHI5*, diploid strain with both *HTB2* alleles tagged with *mCitrine* | This study | 1, S1, S2 |
| KCY028-1 | *Mat α/a; ADE2/ADE2, htb2::HTB2-linker-mCitrine-ADH1term-KlacURA3/htb2::HTB2-linker-mCitrine-ADH1term-CglaTRP1, whi5Δ::CglaTRP1/whi5Δ::kanMX6-LexAprom-WHI5-ADH1term-LEU2, his3/his3::LexA-ER-AD-TF-HIS3* | Inducible *WHI5*, diploid strain with both *HTB2* alleles tagged with *mCitrine* | This study | 1, S1, S2 |
| KCY029-1 | *Mat α/a; ADE2/ADE2, htb2Δ::KlacURA3/htb2::HTB2-linker-mCitrine-ADH1term-CglaTRP1, whi5Δ::CglaTRP1/whi5Δ::kanMX6-LexAprom-WHI5-ADH1term-LEU2, his3/his3::LexA-ER-AD-TF-HIS3* | Inducible *WHI5*, diploid strain with one *HTB2* allele deleted and the other *HTB2* allele tagged with *mCitrine* | This study | 1, S1, S2 |
| KCY031-1 | *Mat α/a; ADE2/ADE2, ura3::HTB1prom-mCitrine-ADH1term-URA3/ura3, WHI5/whi5Δ::kanMX6-LexAprom-WHI5-ADH1term-LEU2, his3/his3::LexA-ER-AD-TF-HIS3* | Inducible *WHI5*, diploid strain with additional copy of *HTB1* promoter expressing *mCitrine* | This study | 3, 5, S6, S7 |
| KCY032-2 | *Mat α/a; ADE2/ADE2, ura3::HHF1prom-mCitrine-ADH1term-URA3/ura3, WHI5/whi5Δ::kanMX6-LexAprom-WHI5-ADH1term-LEU2, his3/his3::LexA-ER-AD-TF-HIS3* | Inducible *WHI5*, diploid strain with additional copy of *HHF1* promoter expressing *mCitrine* | This study | 3, 5, S6, S7 |
| KCY033-2 | *Mat α/a; ADE2/ADE2, ura3::HTB2prom-mCitrine-ADH1term-URA3/ura3, WHI5/whi5Δ::kanMX6-LexAprom-WHI5-ADH1term-LEU2, his3/his3::LexA-ER-AD-TF-HIS3* | Inducible *WHI5*, diploid strain with additional copy of *HTB2* promoter expressing *mCitrine* | This study | 3, S6 |
| KCY035-3 | *Mat α/a; ADE2/ADE2, ACT1prom-mCitrine-ADH1term-CglaTRP1-URA3/ura3, WHI5/whi5Δ::kanMX6-LexAprom-WHI5-ADH1term-LEU2, his3/his3::LexA-ER-AD-TF-HIS3* | Inducible *WHI5*, diploid strain with additional copy of *ACT1* promoter expressing *mCitrine* | This study | 3, S6 |
| KCY038-1 | *Mat α/a; ADE2/ADE2, ura3::300bpHTB1prom-mCitrine-ADH1term-CglaTRP1-URA3/ura3, leu2/LEU2* | Non-inducible *WHI5*, diploid strain with additional 300 bp of *HTB1* promoter | This study | 5, S7 |

| Name | Genotype | Description | Origin | Fig. |
|---|---|---|---|---|
| | | (truncated 5' – 3') expressing *mCitrine* | | |
| KCY039-1 | *Mat α/a; ADE2/ADE2, ura3::300bpHTB1prom-mCitrine-ADH1term-CglaTRP1-URA3/ura3, WHI5/whi5Δ::kanMX6-LexAprom-WHI5-ADH1term-LEU2, his3/his3::LexA-ER-AD-TF-HIS3* | Inducible *WHI5*, diploid strain with additional 300 bp of *HTB1* promoter (truncated 5' – 3') expressing *mCitrine* | This study | 5, S7 |
| KCY040-1 | *Mat α/a; ADE2/ADE2, ura3::450bpHTB1prom-mCitrine-ADH1term-CglaTRP1-URA3/ura3, leu2/LEU2* | Non-inducible *WHI5*, diploid strain with additional 450 bp of *HTB1* promoter (truncated 5' – 3') expressing *mCitrine* | This study | 5, S7 |
| KCY041-1 | *Mat α/a; ADE2/ADE2, ura3::450bpHTB1prom-mCitrine-ADH1term-CglaTRP1-URA3/ura3, WHI5/whi5Δ::kanMX6-LexAprom-WHI5-ADH1term-LEU2, his3/his3::LexA-ER-AD-TF-HIS3* | Inducible *WHI5*, diploid strain with additional 450 bp of *HTB1* promoter (truncated 5' – 3') expressing *mCitrine* | This study | 5, S7 |
| KCY043-1 | *Mat α/a; ADE2/ADE2, ura3::300bpHHF1prom-mCitrine-ADH1term-CglaTRP1-URA3ura3, WHI5/whi5Δ::kanMX6-LexAprom-WHI5-ADH1term-LEU2, his3/his3Δ::LexA-ER-AD-TF-HIS3* | Inducible *WHI5*, diploid strain with additional 300 bp of *HHF1* promoter (truncated 5' – 3') expressing *mCitrine* | This study | 5, S7 |
| KCY045-1 | *Mat α/a; ADE2/ADE2, ura3::450bpHHF1prom-mCitrine-ADH1term-CglaTRP1-URA3/ura3, WHI5/whi5Δ::kanMX6-LexAprom-WHI5-ADH1term-LEU2, his3/his3::LexA-ER-AD-TF-HIS3* | Inducible *WHI5*, diploid strain with additional 450 bp of *HHF1* promoter (truncated 5' – 3') expressing *mCitrine* | This study | 5, S7 |
| KSY212-2 | *Mat α; ADE2, rrp6Δ::CglaTRP1* | Non-inducible *WHI5*, haploid *rrp6Δ* strain | This study | 2, S5 |
| KSY213-6 | *Mat a; ADE2, rrp6Δ::CglaTRP1, whi5Δ::kanMX6-LexAprom-WHI5-ADH1term-LEU2, his3::LexA-ER-AD-TF-HIS3* | Inducible *WHI5*, haploid *rrp6Δ* strain | This study | 2, S5 |
| KSY214-1 | *Mat α; ADE2, hir1Δ::CglaTRP1* | Non-inducible *WHI5*, haploid *hir1Δ* strain | This study | 2, S5 |
| KSY215-2 | *Mat a; ADE2, hir1Δ::CglaTRP1, whi5Δ::kanMX6-LexAprom-WHI5-ADH1term-LEU2, his3::LexA-ER-AD-TF-HIS3* | Inducible *WHI5*, haploid *hir1Δ* strain | This study | 2, S5 |
| KSY219-3 | *Mat a; ADE2, rtt106Δ::CglaTRP1, whi5Δ::kanMX6-LexAprom-WHI5-ADH1term-LEU2, his3::LexA-ER-AD-TF-HIS3* | Inducible *WHI5*, haploid *rtt106Δ* strain | This study | 2, S5 |
| KSY208-3 | *Mat α; ADE2, ura3::mCitrine-ADH1term-URA3* | Non-inducible *WHI5*, haploid strain with additional *mCitrine* copy (not expressed) | This study | |

| Name | Genotype | Description | Origin | Fig. |
|---|---|---|---|---|
| KSY222-1 | *Mat a; ADE2, ura3::HTB1prom-mCitrine-ADH1term-CglaTRP1-URA3, whi5Δ::kanMX6-LexAprom-WHI5-ADH1term-LEU2, his3::LexA-ER-AD-TF-HIS3* | Inducible *WHI5*, haploid strain with additional copy of *HTB1* promoter expressing *mCitrine* | This study | 3, S6 |
| KSY223-3 | *Mat a; ADE2, ura3::HHF1prom-mCitrine-ADH1term-CglaTRP1-URA3, whi5Δ::kanMX6-LexAprom-WHI5-ADH1term-LEU2, his3::LexA-ER-AD-TF-HIS3* | Inducible *WHI5*, haploid strain with additional copy of *HHF1* promoter expressing *mCitrine* | This study | 3, S6 |
| KSY225-2 | *Mat a; ADE2, ura3::HTB2prom-mCitrine-ADH1term-CglaTRP1-URA3, whi5Δ::kanMX6-LexAprom-WHI5-ADH1term-LEU2, his3::LexA-ER-AD-TF-HIS3* | Inducible *WHI5*, haploid strain with additional copy of *HTB2* promoter expressing *mCitrine* | This study | 3, S6 |
| KSY226-3 | *Mat a; ADE2, ura3::mCitrine-ADH1term-URA3, whi5Δ::kanMX6-LexAprom-WHI5-ADH1term-LEU2, his3::LexA-ER-AD-TF-HIS3* | Inducible *WHI5*, haploid strain with additional *mCitrine* copy (not expressed) | This study | |
| KSY229-1 | *Mat α; ADE2, ura3::ACT1prom-mCitrine-ADH1term-CglaTRP1-URA3* | Non-inducible *WHI5*, haploid strain with additional copy of *ACT1* promoter expressing *mCitrine* | This study | 3, S6 |
| KSY230-1 | *Mat a; ADE2, ura3::ACT1prom-mCitrine-ADH1term-CglaTRP1-URA3, whi5Δ::kanMX6-LexAprom-WHI5-ADH1term-LEU2, his3::LexA-ER-AD-TF-HIS3* | Inducible *WHI5*, haploid strain with additional copy of *ACT1* promoter expressing *mCitrine* | This study | 3, S6 |
| MMY116-2C | *Mat α; ADE2* | Non-inducible *WHI5*, haploid strain | Skotheim lab stock | 1, 2, 3, S1, S3, S4, S5 |
| MS62-1 | *Mat a; ADE2, whi5Δ::kanMX6, his3::LexA-ER-AD-TF-HIS3* | β-estradiol dependent transcription factor, haploid *whi5Δ* strain | Matthew Swaffer, Skotheim lab | S3 |
| MS63-1 | *Mat a; ADE2, whi5Δ::kanMX6-LexAprom-WHI5-ADH1term-LEU2, his3::LexA-ER-AD-TF-HIS3* | Inducible *WHI5*, haploid strain | Matthew Swaffer, Skotheim lab | 1, 2, 3, S1, S3, S4, S5 |

**Supplementary Table 1.** Yeast strains used in this work. All strains are based on W303. *CglaTRP1* denotes the *TRP1* gene of the organism *C. glabrata*, *KlacURA3* denotes the *URA3* gene of the organism *K. lactis*.

| qPCR primer | **MS63-1** **[mean $C_p^{Gene}$]** | **Δ strain** **[$Cp$ range]** |
|---|---|---|
| HHO1 | $20.3 \pm 0.1$ | $no\ amp$ |
| HTB2 | $16.8 \pm 0.1$ | $40.9 - no\ amp$ |
| HHF1 | $17.7 \pm 0.3$ | $34.6\ -\ no\ amp$ |
| HHF2 | $19.5 \pm 0.1$ | $33.6 - 35.9$ |
| HHT1 | $18.0 \pm 0.1$ | $32.9 - no\ amp$ |
| HHT2 | $19.0 \pm 1.4$ | $33.7 - no\ amp$ |

**Supplementary Table 2.** Results of qPCR measurements on deletion strains to test for primer specificity. $C_p^{Gene}$ for the MS63-1 strain are the mean of n = 3 technical replicates, with standard deviation. For the deletion strains, $C_p$-ranges reach from the minimum $C_p^{min}$ to maximum $C_p^{max}$ values of n = 3 technical replicates for *HHO1*, *HTB2*, *HHT2* and n = 6 technical replicates for *HHF1*, *HHF2*, *HHT1*. No detectable amplification curve over threshold is denoted as "$no\ amp$".

| Gene | qPCR primer direction | qPCR primer sequence (5’ - 3’) |
|---|---|---|
| *ACT1* | forward | AGTTGCCCCAGAAGAACACC |
| | reverse | GGACAAAACGGCTTGGATGG |
| *ENO2* | forward | TTGTTCCATCTGGTGCCTCC |
| | reverse | ACGAAAGCAGCAGCAATGAC |
| *HHF1* | forward | TACACCGAACACGCCAAGAG |
| | reverse | TTGCTTGTTGTTACCGTTTTCTT |
| *HHF2* | forward | ACGAAGAAGTCAGAGCCGTC |
| | reverse | ACCGATTGTTTAACCACCGATTG |
| *HHO1* | forward | ACCAGCAAAGGCAAGGAGAA |
| | reverse | AAAGCCGTGAGCCCTTCAAT |
| *HHT1* | forward | CAATCTTCTGCCATCGGTGC |
| | reverse | ACTGATGACAATCAACAAACTATGA |
| *HHT2* | forward | AGCAAACACTCCACAATGGC |
| | reverse | CAAGGCAACAGTACCTGGCT |
| *HTA1* | forward | GTTGCCAAAGAAGTCTGCCA |
| | reverse | CAGTTTAGTTCCTTCCGCCTT |
| *HTA2* | forward | TCGCCCAAGGTGGTGTTTT |
| | reverse | TGATTTGCTTTGTTTCTTTTCAACT |
| *HTB1* | forward | TACACACATACAATGTCTGCTAAAG |
| | reverse | AGTGTCAGGGTGAGTTTGCTT |
| *HTB2* | forward | CCTCTGCCGCCGAAAAGAAA |
| | reverse | TCTTACCATCGACGGAGGTTG |
| *mCitrine* | forward | GAGCTGAAGGGCATCGACTT |
| | reverse | TTCTGCTTGTCGGCCATGAT |
| *RDN18* | forward | AACTCACCAGGTCCAGACACAATAAGG |
| | reverse | AAGGTCTCGTTCGTTATCGCAATTAAGC |
| *RPB1* | forward | CCAGAAGTGGTCACACCATATAA |
| | reverse | GGTCTCCGCTATCACGAATG |
| *RPB3* | forward | TGTGGGGTCTATTCCCGTTG |
| | reverse | CGCCCGTCATCATTACGTCT |

**Supplementary Table 3.** Sequences of qPCR primer used in this work.
